# Characterisation of prostate cancer sialome re-engineering via *CMAH* transfection reveals a bystander effect that propagates Neu5Gc presentation to neighbouring cells

**DOI:** 10.64898/2026.09.09.750340

**Authors:** Yumiko Uno, Amanda R. Noble, Esme Hutton, Nathalie Signoret, Martin A. Fascione

**Affiliations:** Department of Chemistry, University of York, York, YO10 5DD, UK; Hull York Medical School, University of York, York, YO10 5DD, UK; Department of Biology, University of York, York, YO10 5DD, UK

**Keywords:** Neu5Gc, Neu5Ac, CMAH, Prostate cancer, LNCaP

## Abstract

Sialic acids are a family of nine-carbon α-keto sugars that play essential roles in human health and disease. In mammals, they are found in two main forms: N-acetylneuraminic acid (Neu5Ac) and N-glycolylneuraminic acid (Neu5Gc), with interactions between Neu5Ac-containing glycans and Siglec receptors on immune cells increasingly recognised as glyco-immune checkpoints, promoting immunosuppression. However, humans do not synthesise Neu5Gc due pseudogenisation of the *CMAH* gene which encodes the cytidine monophospho-N-acetylneuraminic acid hydroxylase enzyme responsible for CMP-Neu5Ac conversion into CMP-Neu5Gc, which then serves as the donor substrate for sialyltransferases. Here, we investigated the effects of re-engineering tumour cell-surface glycans in a prostate cancer cell model by expressing rat *CMAH*, thereby enabling the conversion of CMP-Neu5Ac to CMP-Neu5Gc. LNCaP cells transfected with the rat *CMAH* gene predominantly incorporated Neu5Gc into mucin-associated O-glycans implicated in immune suppression. Treatment with sialidase significantly reduced Neu5Gc expression, indicating that Neu5Gc was presented on cell-surface glycans, while cell-tracing experiments demonstrated the transfer of Neu5Gc to neighbouring cells, revealing a potential bystander effect capable of propagating Neu5Gc expression within the tumour microenvironment.

## Introduction

Sialic acid is the general term often used to describe 9-carbon α-keto acid sugars, a diverse class of biological molecules abundant on the surface of animal cells^1^ and microbes^2^, with the most prominent members including the ubiquitous *N-*acetylneuraminic acid (Neu5Ac) and *N-*glycolylneuraminic acid (Neu5Gc)^3^. The biological importance of Neu5Ac to human physiology is reflected in its abundance on the outer cell membrane, the interior of lysosomal membranes, as well as on secreted glycoproteins and richly glycosylated mucins. This distribution suggests a role in stabilizing molecules and membranes while regulating interactions with the surrounding environment^4^. However, changes in sialylated glycan expression are also frequently seen during the progression of cells to malignancy and tumour formation^5^, with mucin sialylation the subject of fervent attention in recent years^6^. Specifically, it is now well established that cell-surface sialylated mucins can interact with sialic acid-binding immunoglobulin-like lectins (Siglecs) expressed on immune cells, resulting in immune suppression and facilitating immune evasion by hypersialylated tumour cells^7^. This reveals cell surface sialylation as a potential Achilles’ heel for targeting in new glyco-immunotherapy approaches for treating cancer^8,9^. This therapeutic potential extends to prostate cancer (PCa), which is the second most common cancer and the fifth most aggressive malignancy among men worldwide^10^. Notably, ligands for Siglec-7 and Siglec-9 are abundantly expressed on PCa cells and in human prostate tumour specimens^11^, and blocking Siglec-7/9–sialic acid interactions has been shown to inhibit tumour growth and enhance immune cell infiltration in PCa xenograft models using humanised mice^12^. The biosynthesis of these sialylated ligands is governed by sialyltransferase enzymes (STs), which can be upregulated in PCa^11,13–15^, and use the activated sugar nucleotide derivative cytidine monophosphate-N-acetylneuraminic acid (CMP-Neu5Ac) as a substrate to construct sialylglycoconjugates (Fig. 1a). Interestingly, Neu5Gc, which is a non-human sialic acid, has also been detected in human tumours including PCa^16^. Although widespread among other mammals^17^, humans lack Neu5Gc biosynthesis capability due to a 92-bp deletion in the *CMAH* gene^18^ coding for the enzyme cytidine monophospho-N-acetylneuraminic acid hydroxylase which catalyses the transfer of an oxygen atom to CMP-Neu5Ac to generate CMP-Neu5Gc (Fig. 1b)^19^. It is now understood that Neu5Gc presentation on human cells results from the metabolic incorporation of this sialic acid from dietary sources^20,21^. Here, Neu5Gc enters the cell through pinocytosis^22^ (Fig. 1a) before hijacking the latter stages of the sialylation pathway, first through activation by the CMP synthetase enzyme (CMAS) and subsequently ST-mediated sialoglycoconjugate synthesis. Despite the subtle chemical difference between Neu5Ac and Neu5Gc, this reengineering of the host cell sialome can profoundly impact how Neu5Gc-presenting cells interact with their microenvironment. This includes tuning binding to Siglecs^23^ and complement factor H^24,25^, and altering interactions with pathogens such as influenza^26,27^ and malaria-causing protozoa^28^. Furthermore, with evidence that humans possess circulating anti-Neu5Gc antibodies^29,30^, interactions between these antibodies and metabolically incorporated Neu5Gc may promote chronic inflammation, potentially driving tumour progression^31^. As such, it is essential to establish robust experimental models to study Neu5Ac-to-Neu5Gc reengineering and investigate its impact on cancer cells. Herein, we characterise the extent and nature of the sialome reengineering achieved in prostate cancer cells through *CMAH* transfection, and explore whether shedding and subsequent uptake of Neu5Gc-bearing molecules by neighbouring non-CMAH-expressing cells, may propagate this reengineering in a tumour microenvironment.

**Figure 1.**
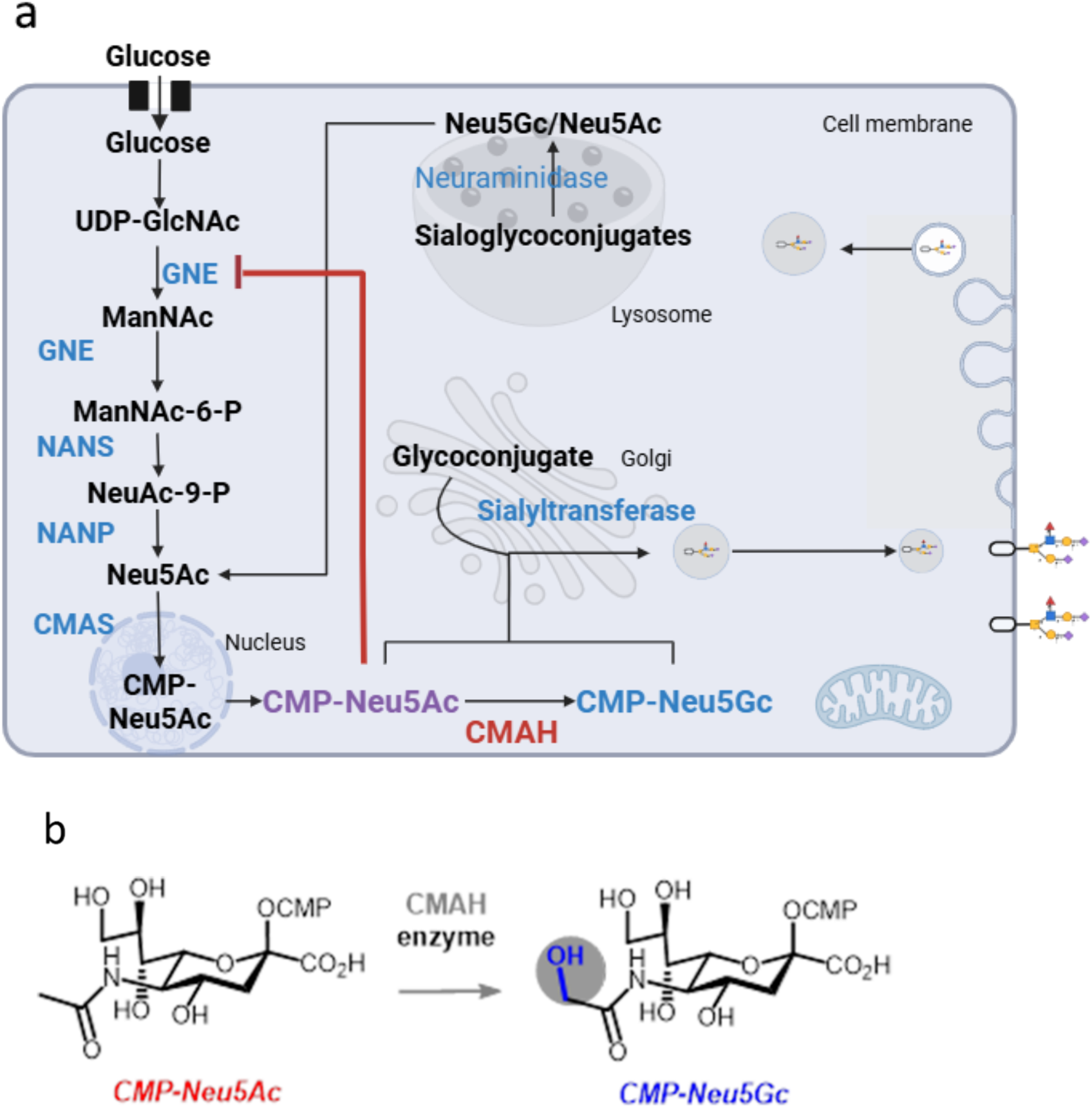
Schematic overview of the sialic acid metabolic pathway in humans. (a) Sialic acid biosynthesis is initiated by the bifunctional enzyme GNE, which plays a central regulatory role^32^. Subsequently synthesised Neu5Ac is transported from the cytosol into the nucleus, where CMAS catalyses the addition of a CMP group to form the activated compound CMP-Neu5Ac before transfer back to the cytosol^33^. CMP-Neu5Gc can then be constructed in the cytosol if CMAH is present (normally absent in humans), which adds an oxygen atom to the N-acetyl group of CMP-Neu5Ac. Both CMP-Neu5Ac and CMP-Neu5Gc can then be transported into the Golgi apparatus by the CMP-sialic acid transporter^34^. Inside the Golgi, sialyltransferases facilitate the transfer of sialic acids from their CMP-activated forms to the terminal positions of glycoconjugates, thereby completing the sialylation process^35^. Regulation of this pathway is achieved in part through feedback inhibition of GNE by CMP-Neu5Ac, as illustrated in the schematic (red line). Sialic acid and glycoconjugates can also enter human cells through pinocytosis^22^, before sialic acids can be released and recycled by lysosomal sialidase and subsequently transported into the cytosol via the lysosomal sialic acid transporter, where they are available for activation and incorporation into glycoconjugates. (b) Biosynthesis of CMP-Neu5Gc from CMP-Neu5Ac through the action of the CMAH hydroxylase enzyme in the cytosol.

## Results

We chose to use Lymph Node Carcinoma of the Prostate (LNCaP)^36^ as a model cell line for our experiments, as it is among the most extensively studied PCa cell lines, is androgen-dependent^37^ and exhibits relatively slow growth and indolent biological behaviour, reflecting the characteristics of most clinically observed prostate cancers^36^. We first sought to select a functional CMAH enzyme for expression on LNCaP cells by comparing the amino acid sequences of CMAH proteins from different mammalian species (Fig. 2). A study by Song *et al.* identified several putative functional regions in the CMAH amino acid sequence^38^, including a CMP-Neu5Ac-binding site (red box), a Rieske iron-sulfur cluster binding site (blue box), a mononuclear iron-binding site (green boxes), and a cytochrome b5 interaction site (yellow box). The putative CMP-Neu5Ac-binding site was assigned through sequence comparison with conserved CMP-Neu5Ac-binding motifs in three sialyltransferases^39^. Although there is a negligible sequence difference between species, we noted that the Neu5Ac-binding site (red box) of the zebrafish CMAH differs subtly from other species. Specifically, histidine (H) and tyrosine (Y) residues are present in place of an otherwise conserved serine (S) and phenylalanine (F) pair. As these substitutions occur within the putative substrate-binding region, they could affect substrate recognition or the catalytic activity of the enzyme. Neu5Gc abundance also varies among species and tissues^40^. This variation may arise from differences in CMAH expression, stability or catalytic activity, together with tissue-specific differences in substrate and cofactor availability and the metabolism of Neu5Gc-containing glycoconjugates. We therefore hypothesised that species-specific differences in the CMAH amino acid sequence might influence Neu5Gc biosynthesis. To investigate this possibility under the same cellular conditions, we compared LNCaP cells transfected with either rat or zebrafish CMAH coding sequences.

**Figure 2.**
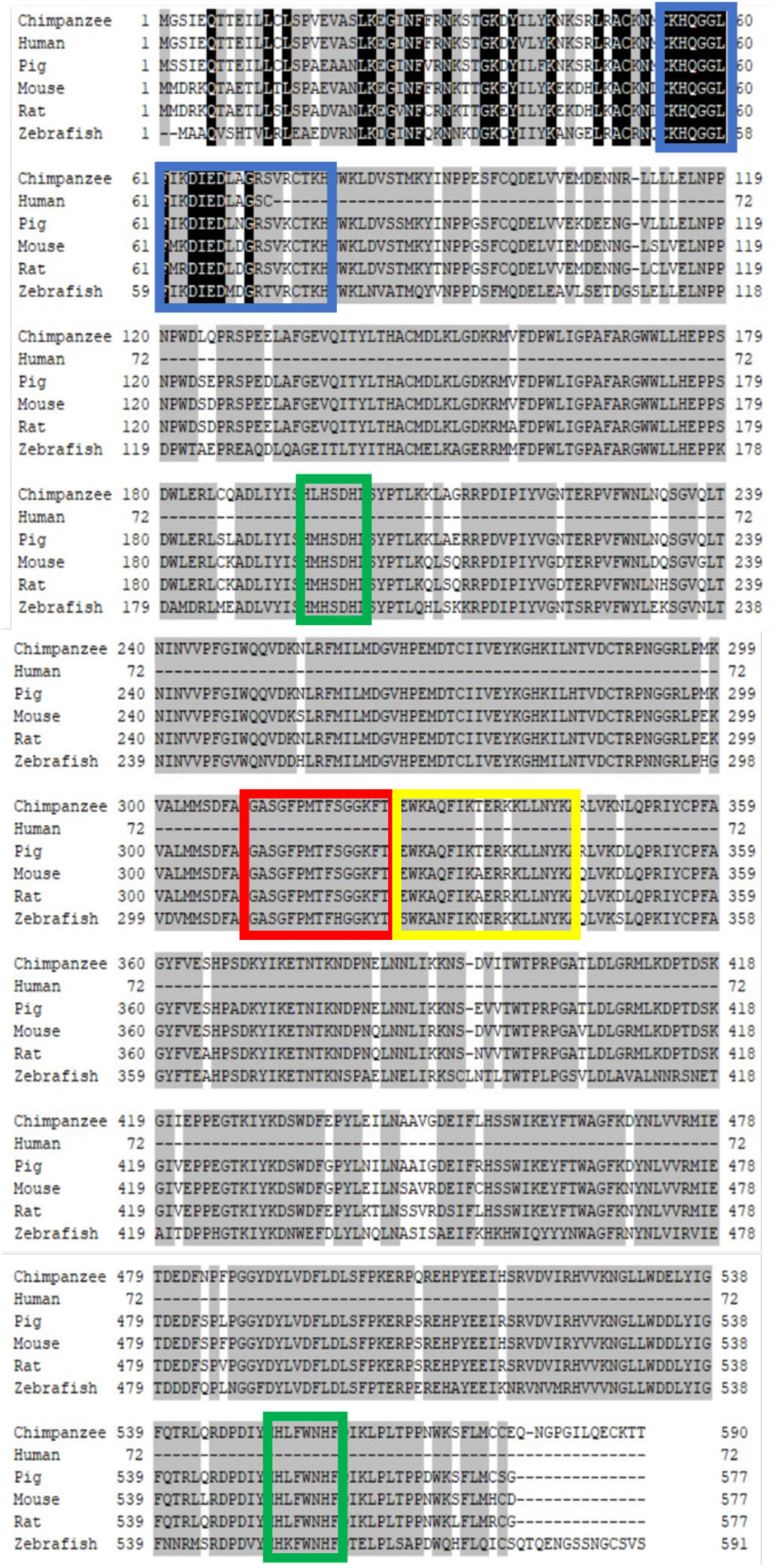
Conserved functional motifs in CMAH proteins across vertebrate species. A multiple-sequence alignment of the deduced CMAH amino acid sequences is shown for chimpanzee (NP_001009041), pig (NP_001106486), mouse (NP_001104580), rat (NP_001019444.1), zebrafish (NP_001002192), and the putative human CMAH sequence (AAC68881). Human *CMAH* is non-functional and encodes a truncated sequence of only 72 amino acids because of a 92-bp frameshifting exon deletion^41^. Residues identical across all six sequences are highlighted in black, whereas residues conserved in five or four sequences are highlighted in dark grey and light grey, respectively. The coloured boxes indicate putative functional regions assigned according to Song *et al.* ^38^: red, the predicted CMP-Neu5Ac-binding site; blue, the Rieske iron–sulfur cluster-binding site; green, the putative mononuclear iron-binding sites; and yellow, the possible cytochrome b₅ interaction site.

To gain further insight into the differences between rat and zebrafish CMAH, we compared their predicted three-dimensional structures using the AlphaFold2 Colab implementation^42^. The two homologues exhibited highly similar overall structures, with only minor differences observed (Fig. 3a–c). Notably, both models contained a cavity encompassing residues proposed to form the CMP-Neu5Ac-binding site (Fig. 3d), and the predicted cavity dimensions appeared compatible with accommodation of the CMP-Neu5Ac substrate. Despite the identified amino acid substitutions, the AlphaFold2 models therefore predicted little difference in the overall three-dimensional organisation of the putative substrate-binding region between rat and zebrafish CMAH.

**Figure 3.**
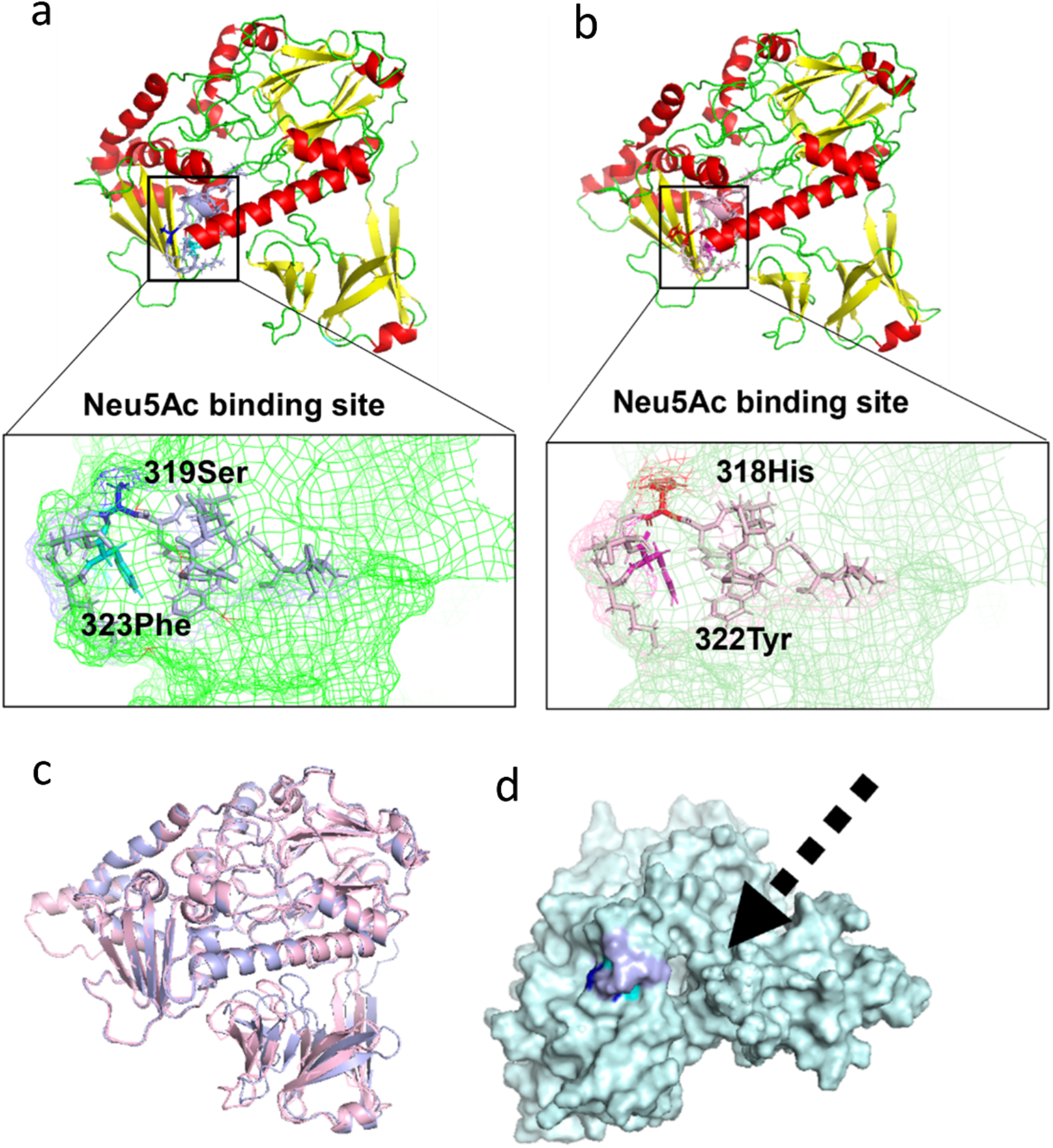
PyMOL analysis of CMAH AlphaFold models. (a,b) Predicted structures of rat (a) and zebrafish (b) CMAH, with α-helices shown in red, β-strands in yellow, and flexible loops in green. Conserved residues are highlighted. Residues associated with the previously proposed CMP-Neu5Ac-binding site are indicated: Ser319 (blue) and Phe323 (cyan) in rat CMAH, and His318 (orange) and Tyr322 (magenta) in zebrafish CMAH. (c) Structural alignment of the rat (light blue) and zebrafish (light pink) CMAH models. (d) Surface representation of the rat CMAH model showing the predicted cavity encompassing the proposed CMP-Neu5Ac-binding site. Ser319 and Phe323 are shown in blue and cyan, respectively, and the arrow indicates the cavity.

To quantify cell-surface Neu5Gc produced by the two CMAH homologues, we used flow cytometry with a polyclonal chicken IgY anti-Neu5Gc antibody^43^. LNCaP cells were transiently transfected with plasmids encoding either functionally active rat or zebrafish *CMAH* cDNA under the control of a CMV promoter. To minimise the impact of Neu5Gc originating from foetal bovine serum (FBS) used for routine cell culture, RPMI containing 10% FBS (R10) was replaced 24 hours post transfection by Neu5Gc-free 5% containing human serum medium (R5-HS), and the LNCaP cells cultured for a further 72 hours to allow time for glycan biosynthesis and remodelling (Fig. 4a). Mean Fluorescence Intensity (MFI) associated with anti-Neu5Gc binding was detected using an AF488-coupled anti-chicken secondary antibody (Fig. 4b). These experiments showed a increase in Neu5Gc detected at the surface of rat or zebrafish *CMAH -*transfected [CMAH (+)] cells compared to non-transfected (WT) controls, with a full histogram shift for AF488 signal indicating that the entire population of transfected cells expressed Neu5Gc on their surface. However no difference was observed between cells expressing either of the two functionally active *CMAH* gene homologs (Fig. 4c).

**Figure 4.**
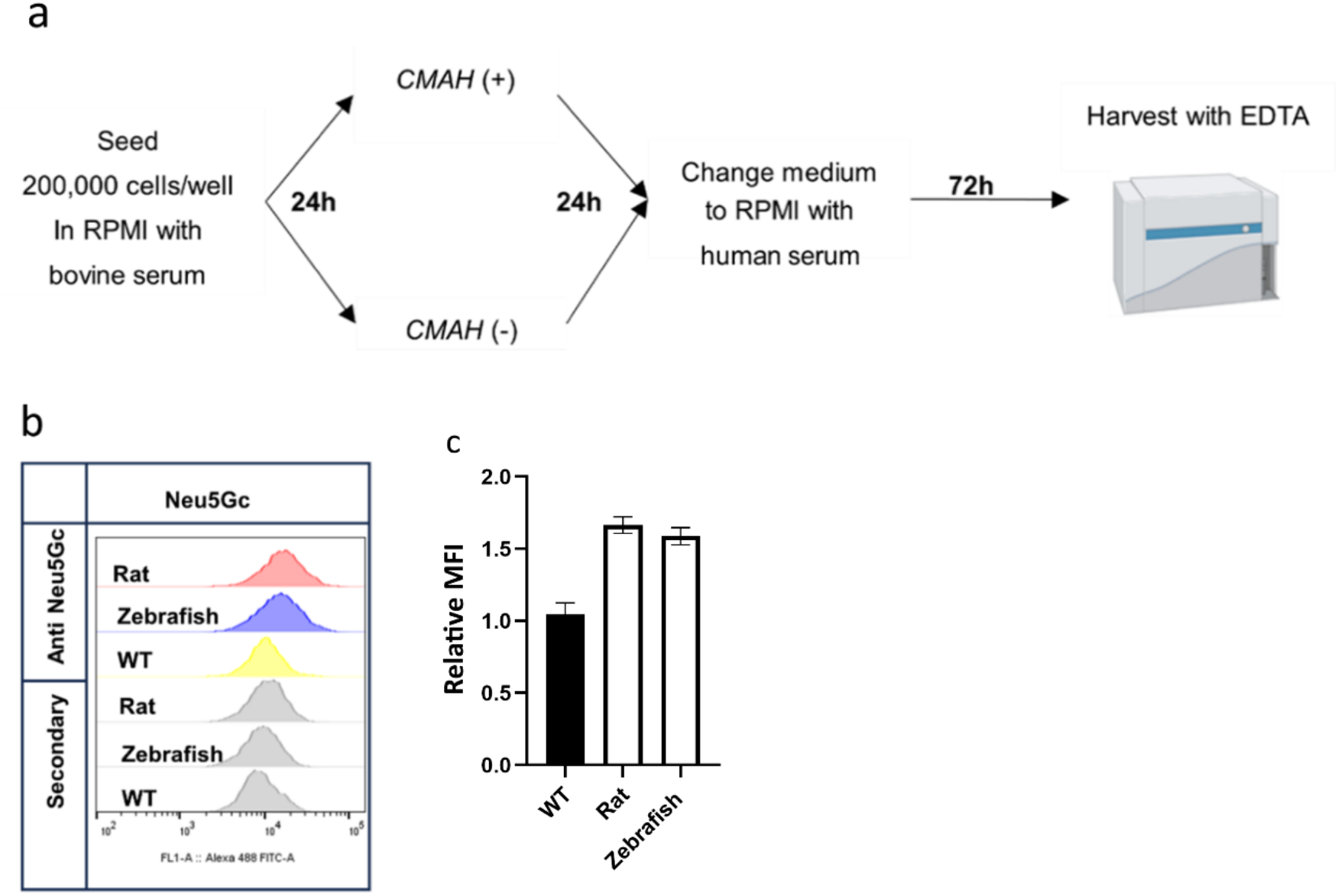
Neu5Gc expression in LNCaP cells transfected with rat and zebrafish *CMAH*. (a) Overview of the experimental workflow. LNCaP cells were cultured for a total of five days. Following transfection on day 2, the standard R10 medium containing 10% FBS was replaced with R5-HS medium containing 5% human serum on day 3. On day 6, Neu5Gc expression was analysed by flow cytometry following antibody staining. (b) Cells were stained with anti-Neu5Gc antibodies, followed by anti-chicken AF488. The histogram overlay shows AF488 (FITC-A) fluorescence intensity in *CMAH* (+) LNCaP cells (rat: red; zebrafish: blue), wild type (WT: yellow), and secondary antibody-only controls for each (grey). (c) The graph presents anti-Neu5Gc mean fluorescence intensity (MFI) values obtained from one experiment performed using three technical replicate wells per condition. MFI values were normalised to those of non-transfected WT cells (WT = 1). Bars represent the mean ± SD of three technical replicates.

Having established that CMAH homologues from different species were functionally comparable in this experimental system, we next investigated the temporal effects of rat CMAH expression on LNCaP cells. Cells were cultured for 72, 96 or 120 h following transient transfection with the rat *CMAH* gene (Fig. 5a). At all three time points, the flow-cytometry histograms showed a rightward shift (Fig. 5b), indicating successful cell-surface presentation of Neu5Gc following transfection. However, the greatest increase in cell-surface Neu5Gc was observed at 96 h, after which the signal declined at 120 h (Fig. 5c). Because cell density and culture duration can affect cell-surface glycan composition^44^, added to the fact that transfection by itself may impact on cell metabolism and growth, Neu5Gc levels in CMAH (+) LNCaP cells were compared at each time point with those in the corresponding WT and secondary-antibody-only controls.

**Figure 5.**
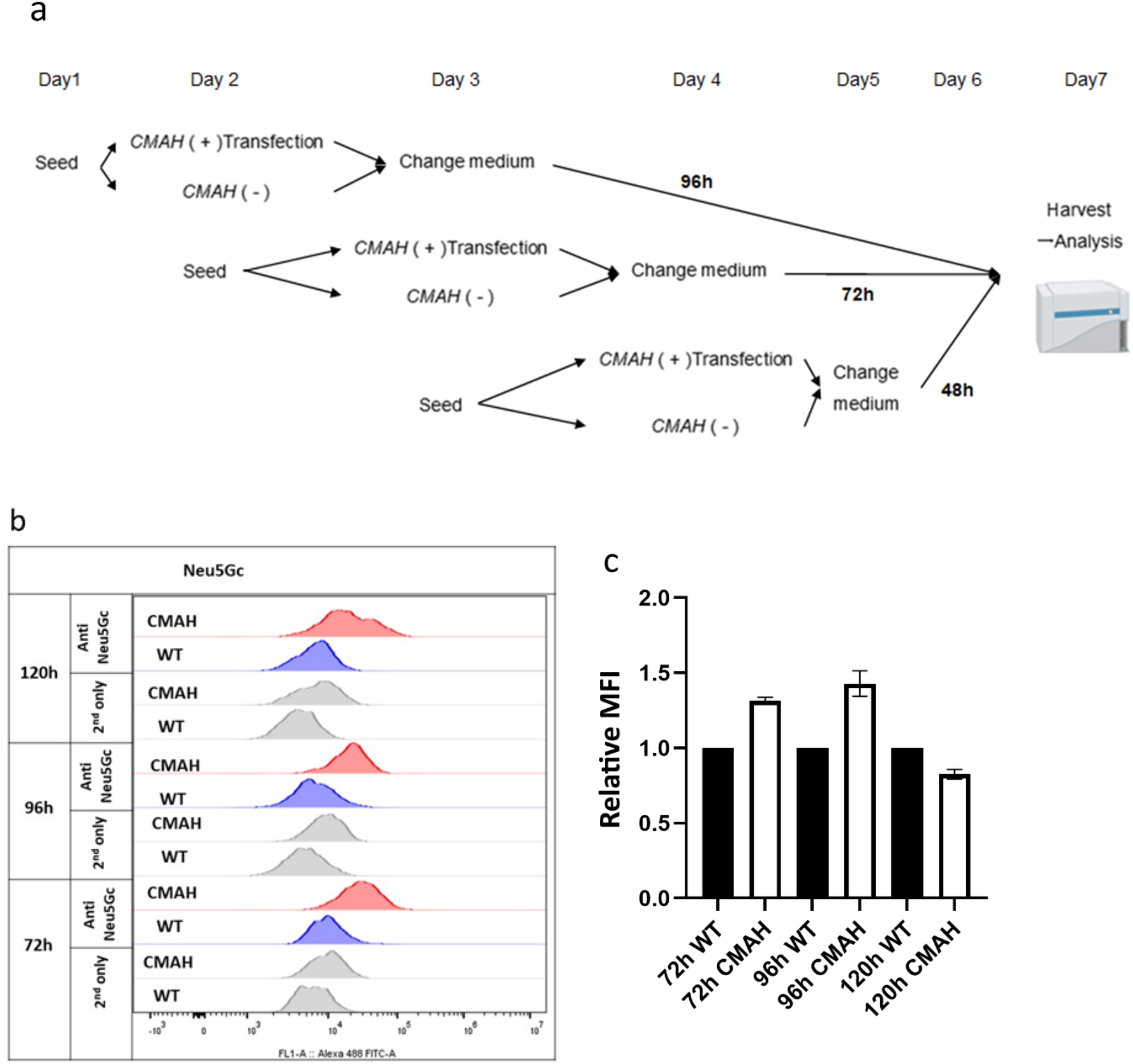
Comparison of transfection effects at 72 h, 96 h, and 120 h post-transfection. (a) Schematic overview of the experimental workflow. LNCaP cells were seeded and transfected on staggered days so that all samples could be harvested and analysed simultaneously on day 7. At 24 h after transfection, the standard R10 medium containing 10% FBS was replaced with R5-HS medium supplemented with 5% human serum. Cells were then cultured for a further 48, 72 or 96 h, corresponding to total post-transfection periods of 72, 96 or 120 h, respectively. Cell-surface Neu5Gc was subsequently analysed by flow cytometry. (b) Histogram overlay showing AF488 fluorescence intensity (FITC-A) at 72 h, 96 h, and 120 h post-transfection. Red: CMAH (+) cells; Blue: WT; Grey: secondary antibody-only controls for both WT and CMAH (+) cells. (c) The graph presents anti-Neu5Gc MFI values obtained from one experiment performed using three technical replicate wells per condition. At each time point, the mean MFI of the corresponding WT control was set to 1, and the CMAH (+) values were normalised accordingly. Bars represent the mean ± SD of three technical replicate wells.

We next examined whether a commercially available neuraminidase could remove Neu5Gc from the surface of CMAH (+) cells. Neu5Gc was detected by flow cytometry using intact cells, allowing its presentation to be assessed in the native cell-surface glycoconjugate context rather than in denatured protein extracts, as would be the case for Western blotting. To confirm that the increase in anti-Neu5Gc binding resulted from cell-surface Neu5Gc presentation, LNCaP cells were treated with neuraminidase from *Vibrio cholerae*, a broad-specificity enzyme that cleaves terminal α2,3-, α2,6,-and α2,8-linked sialic acids from both N-and O-glycans (Fig. 6a). Following harvesting, CMAH (+) cells were incubated with neuraminidase for 1 h at 37 °C. This treatment significantly reduced Neu5Gc detection (P < 0.05; Fig. 6b,c) to a level approaching that observed in WT cells, supporting the conclusion that the detected signal reflected the re-engineering of cell-surface glycans from Neu5Ac to Neu5Gc.

**Figure 6.**
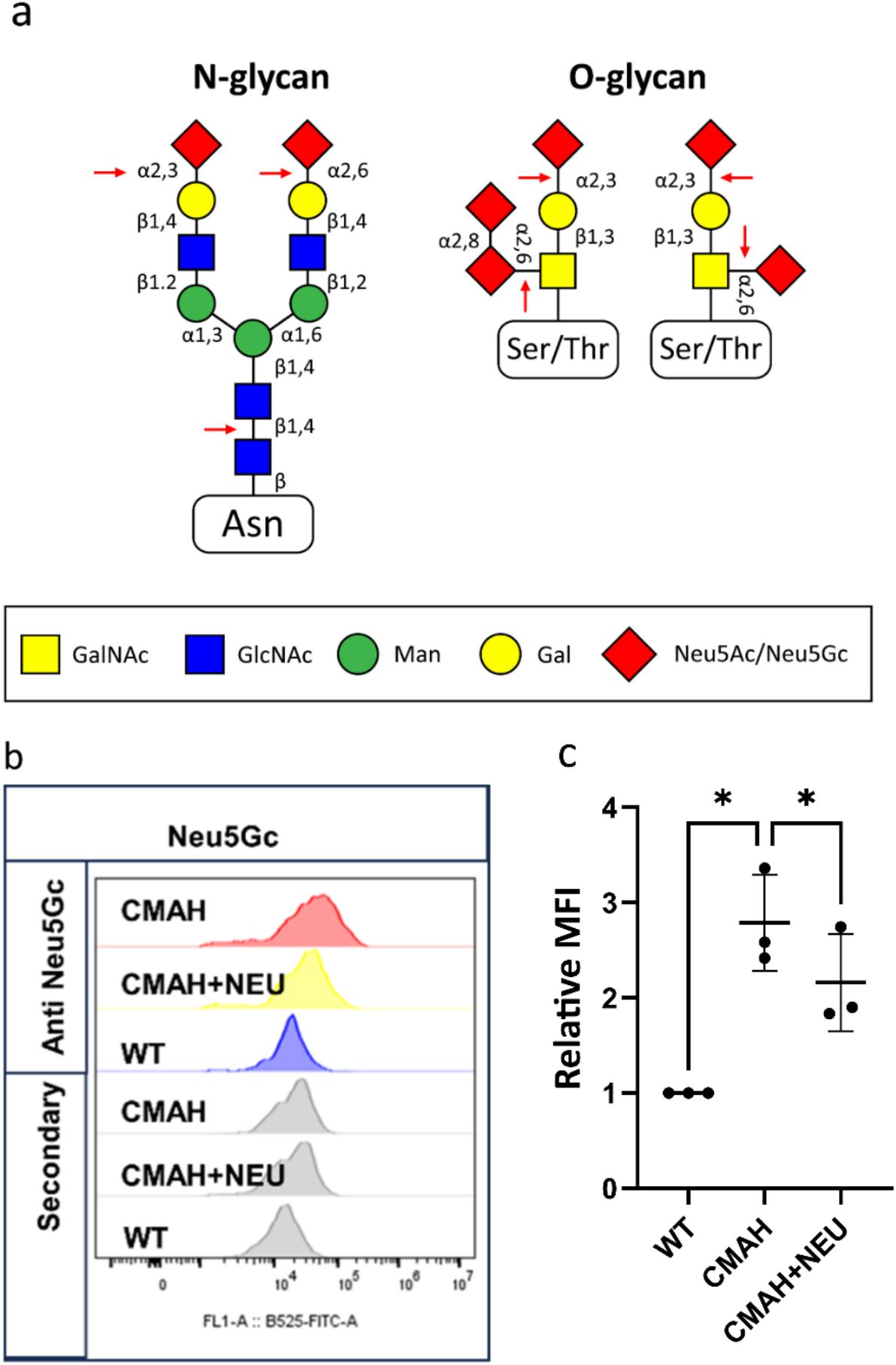
The effect of neuraminidase treatment on CMAH (+) cells. (a) The *V. cholerae* neuraminidase can cleave terminal sialic acid residues by hydrolysing α2,3-, α2,6-, and α2,8-glycosidic linkages in both N-and O-glycans. (b) Cells were stained with anti-Neu5Gc antibodies, followed by AF488 anti-chicken IgY. The histogram overlay shows FITC-A fluorescence intensity for CMAH (+) cells (CMAH, red), CMAH (+) cells treated with neuraminidase (CMAH+NEU, yellow), WT cells (WT, blue), and secondary antibody-only controls for each condition (grey). (c) The graph presents data from three independent experiments. Within each experiment, the MFI was normalised to that of the corresponding WT control (WT = 1). Each point represents the mean of three technical replicate wells from one independent experiment. Data are presented as the mean ± SD (n = 3 independent experiments). CMAH (+) values were compared with the theoretical WT value of 1 using a two-tailed one-sample t-test. Paired CMAH (+) and neuraminidase-treated CMAH (+) values were compared using a two-tailed paired t-test. Holm’s correction was applied across the two comparisons. *:*P* < 0.05

Next we investigated the extent to which N-or O-glycans on the surface of the LNCaP cells had been re-engineered to present Neu5Gc sugars. We used 25 µM 1-deoxymannojirimycin (dMNJ) to inhibit N-glycan processing and 2 mM benzyl-α-GalNAc to inhibit O-glycan biosynthesis^45^, allowing us to dissect the nature of the cell surface Neu5Gc sialylation. As such, 24 hours after *CMAH* transfection, LNCaP cells were incubated with the respective glycosylation inhibitors for 72 hours at 37 °C. Benzyl-α-GalNAc acts as a metabolic decoy and competitive acceptor for downstream glycosyltransferases, thereby disrupting the elongation and sialylation of protein-bound O-glycans^46^. (Fig. 7a), whilst dMNJ modulates N-glycan biosynthesis by inhibiting Golgi α-mannosidase I, an enzyme responsible for trimming high-mannose structures to allow further elongation by N-acetylglucosamine and lactosamine sequences^47^ (Fig. 7b). Subsequently WT cells, CMAH (+) cells, and CMAH (+) cells treated with the N/O-glycan inhibitors were analysed by flow cytometry to again assess the binding of the anti-Neu5Gc antibodies. Notably treatment of CMAH (+) LNCaP cells with benzyl-α-GalNAc significantly reduced anti-Neu5Gc antibody binding (p < 0.01), indicating decreased O-glycan (Neu5Gc) sialylation, whilst in contrast treatment with dMNJ had no significant effect.

**Figure 7.**
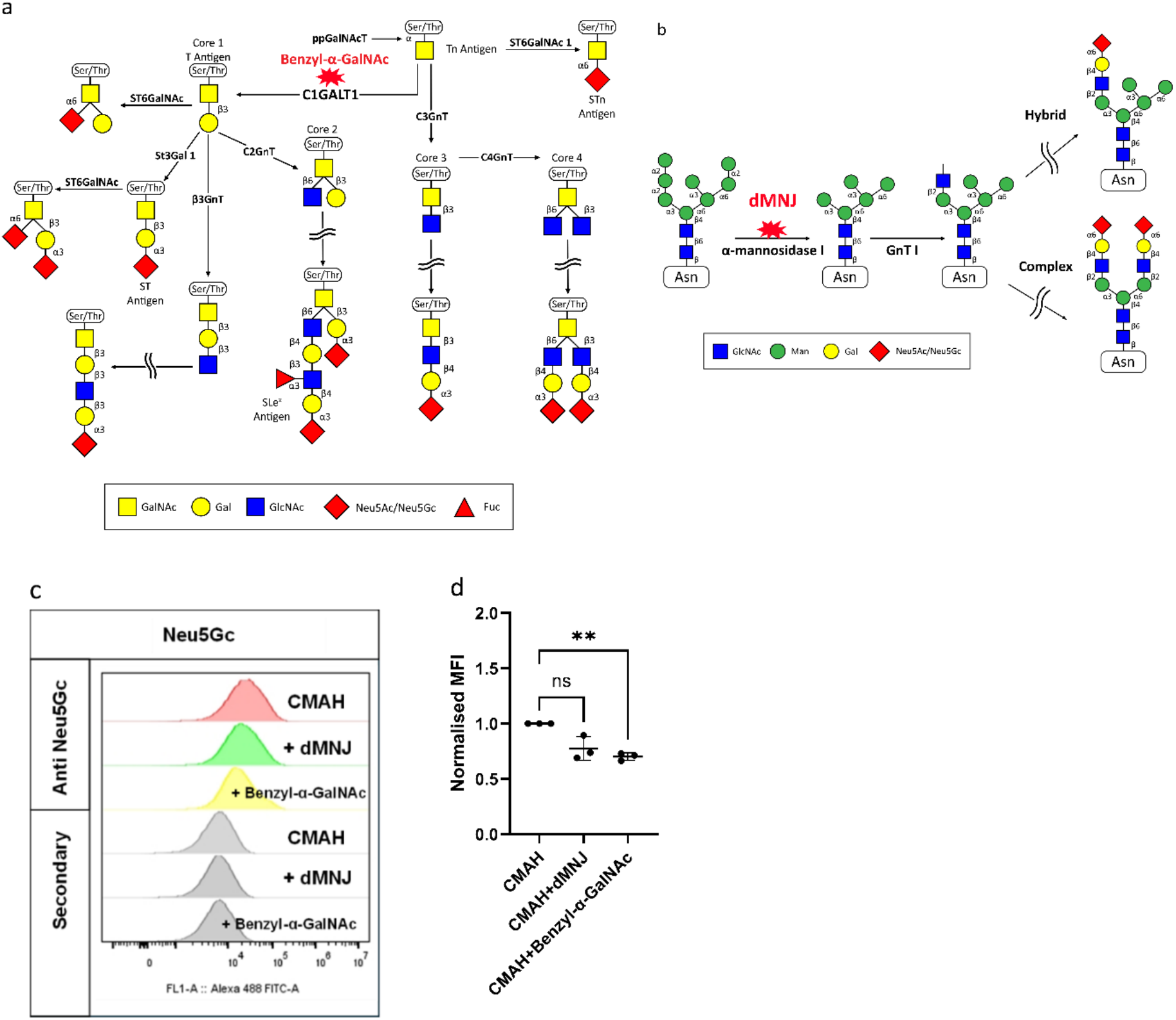
N-and O-glycosylation inhibitor analysis in LNCaP cells. (a) Schematic representation of the O-glycosylation inhibitor Benzyl-α-GalNAc. Benzyl-α-GalNAc acts as a metabolic decoy and competitive acceptor for glycosyltransferases involved in O-glycan extension, thereby disrupting the elongation and terminal sialylation of protein-bound O-glycans. (b) Schematic representation of the N-glycosylation inhibitor dMNJ. dMNJ inhibits Golgi α-mannosidase I, an enzyme involved in trimming high-mannose N-glycans, thus preventing further elongation by N-acetylglucosamine and lactosamine sequences. (c) Cells were stained with anti-Neu5Gc antibodies, followed by AF488 anti-chicken IgY. The histogram overlay shows FITC-A fluorescence intensity for CMAH (+) cells (CMAH, red), CMAH (+) cells treated with dMNJ (+dMNJ, green), CMAH (+) cells treated with Benzyl-α-GalNAc (+Benzyl-α-GalNAc, yellow), and secondary antibody-only controls for each condition (grey). (d) The graph presents anti-Neu5Gc MFI values obtained from three independent experiments. Within each experiment, MFI values were normalised to those of the untreated CMAH(+) control (CMAH(+) = 1). Each point represents the mean of three technical replicate wells from one independent experiment. Data are presented as the mean ± SD (n = 3 independent experiments). Values for each inhibitor-treated condition were compared with the theoretical untreated value of 1 using two-tailed one-sample t-tests. Holm’s correction was applied across the two comparisons. ns: not significant; \*\**P*: < 0.01.

To unpick the cell surface glycan re-engineering in more detail glycopeptide analysis of both CMAH (+) and WT LNCaP cells was conducted. Briefly, cell lysates were digested using an S-Trap-based protocol^48^, which facilitates the use of SDS during sample preparation, and a total of 78 O-glycan and 309 N-glycan structures (Supplementary data), known to occur in mammals, were set as variable modifications. This approach yielded the identification of 2,789 proteins in WT cells and 3,597 proteins in CMAH (+) cells. Ranked intensity plots of the identified proteins and scatterplots of precursor m/z against precursor mass error are shown in Supplementary Figures S1-4. For non-transfected WT cells, 257 unique glycopeptides were identified. Among them, 29 glycopeptides contained Neu5Ac in their glycan structures (Supplementary Table S1). Several well-characterised glycoproteins bearing Neu5Ac-containing glycans were identified, and all sialylated glycoproteins detected in the WT were O-glycans (Fig. 8c), with the peptide profiles of LNCaP cells in this study consistent with those reported by Shah^49^. We focused particularly on Neu5Ac-containing glycoproteins annotated as being localised to the cell surface. Among these, transferrin receptor protein 1 (UniProt accession P02786; TFR1_HUMAN), a widely expressed cell-surface glycoprotein, was identified as carrying a sialylated glycopeptide. As anticipated, no convincing Neu5Gc-containing glycopeptides were detected in WT cells following manual inspection of the spectra. By contrast, for CMAH (+) cells there were 406 glycopeptides matched, with 42 glycopeptides containing Neu5Ac and five glycopeptides containing Neu5Gc in their glycan structure identified (Supplementary Table S2), the majority being O-glycans (Fig. 8a-d). The subcellular localisation of the identified proteins was then assessed using the online database resource The Human Protein Atlas^50^, and although 15 proteins (HNRNPU, KRT10, HSPA5, PDIA3, FOLH1, C1QBP, PHB2, GOT2, ENO1, P4HB, HSP90AB1, CALR, PHB, HSPD1, and ATP5B) among the 100 highest intensity proteins were annotated as cell surface-associated, none of these were sialylated. Since flow cytometry showed Neu5Gc-containing glycans on the cell surface following *CMAH* transfection, we therefore considered whether our glycopeptide analysis was providing a complete picture of the glycoproteins that had been remodelled.

**Figure 8.**
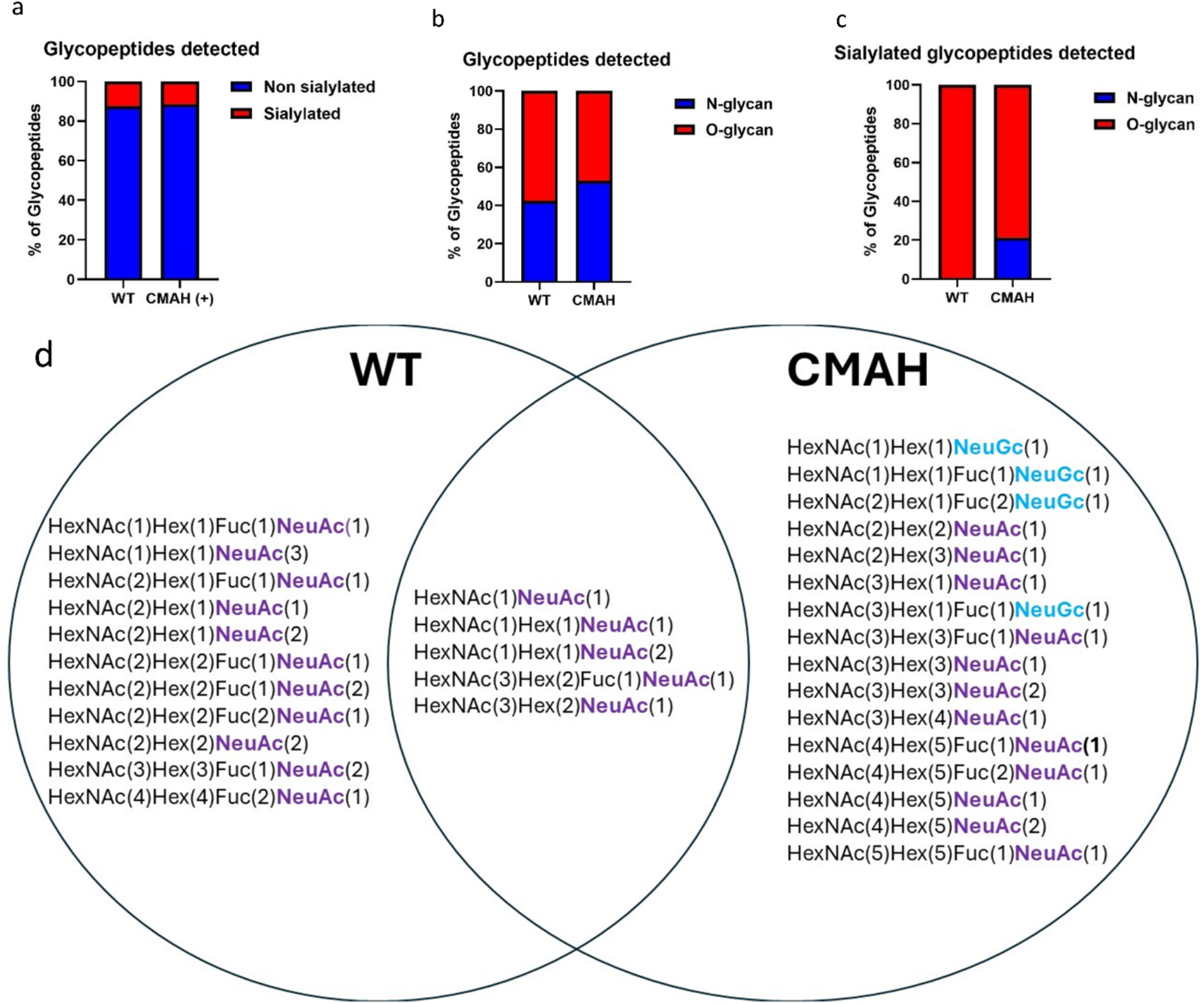
Glycopeptide analysis of WT and CMAH (+) LNCaP cells. (a) Proportion of sialylated glycopeptides (red) relative to non-sialylated glycopeptides (blue) among all detected glycopeptides in WT (left) and CMAH (+) (right) LNCaP cells. (b) Proportion of N-glycans (blue) and O-glycans (red) among all detected glycopeptides in WT (left) and CMAH (+) (right) LNCaP cells. (c) Proportion of N-glycans (blue) and O-glycans (red) among all sialylated glycopeptides in WT (left) and CMAH (+) (right) LNCaP cells. (d) Venn diagram illustrating the overlap and distribution of sialylated glycopeptides detected in WT (left circle) and CMAH (+) (right circle) LNCaP cells.

Notably, no cell surface mucins, which are densely O-glycosylated, were detected in our glycopeptide analysis, despite previous work on transmembrane glycoprotein mucin 1 (MUC1)^51^ indicating that MUC1 overexpression occurred in 58% of primary prostate cancers and 90% of lymph node metastases, while no expression was detected in normal adult or benign tissues^52^. We suspected this may be a result of the established limited cleavage of glycosylated mucins by the trypsin protease used in the S-Trap proteomics protocol^53,54^. In light of the well appreciated challenges of performing mucinomics^55^, we therefore opted to characterise mucin sialylation re-engineering with flow cytometry using mucin-specific proteases. This included the mucinase StcE, derived from an enterohemorrhagic *Escherichia coli* strain, which selectively recognises and cleaves heavily glycosylated mucin domains but does not act on non-mucin O-glycoproteins^56^ (Fig. 9a), and the immunomodulating metalloprotease (IMPa) a broad specificity O-glycoprotease from *Pseudomonas aeruginosa*^57^, which cleaves immediately N-terminal to O-glycosylated serine or threonine residues^58^ (Fig. 9a). Although flow cytometric analysis following mucinase treatment proved technically challenging, as mucinases reduced the number of viable cells, we were able to demonstrate when comparing LNCaP WT cells, CMAH (+) *cells, and* CMAH (+) cells treated with mucinases StcE and IMPa, that Neu5Gc levels on CMAH (+) cells were indeed significantly reduced after treatment with both mucinases (P ≤ 0.01, Fig. 9b and c).

**Figure 9.**
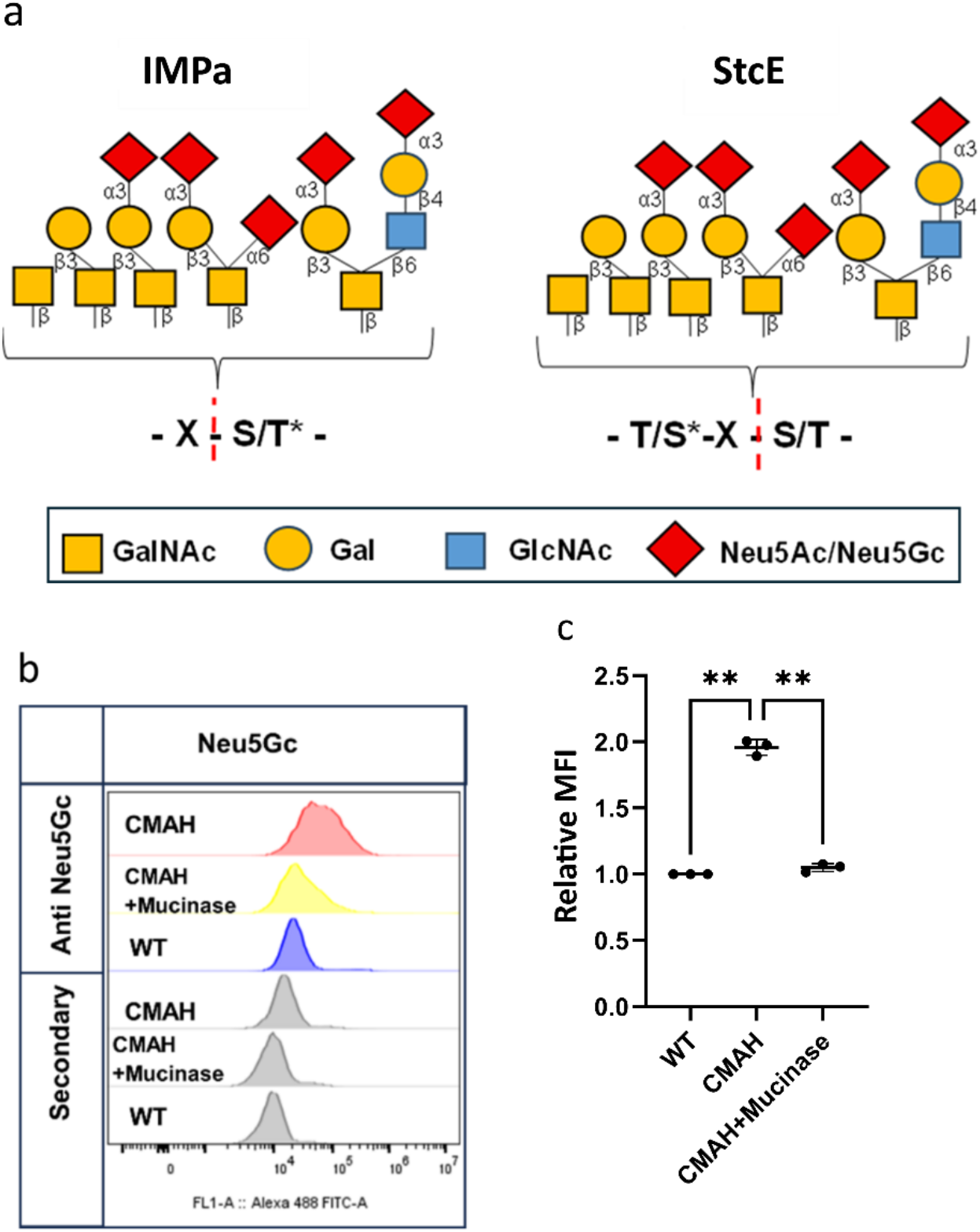
Mucinase analysis in WT and CMAH (+) LNCaP cells. (a) Cleavage motifs for mucinases and O-glycoproteases as described in the literature. The accepted subsite nomenclature for amino acid residues and glycan moieties of glycoprotein substrates is shown, with IMPa and StcE used as representative examples. Red dashed lines indicate the enzymatic cleavage sites. (b) Cells were stained with the anti-Neu5Gc antibodies, followed by AF488 anti-chicken IgY. The histogram overlay shows FITC-A fluorescence intensity for CMAH (+) cells (CMAH, red), CMAH (+) cells treated with mucinase (CMAH + Mucinase (IMPa and StcE), yellow), wild-type cells (WT, blue), and secondary antibody-only controls for each condition (grey). (c) The graph presents anti-Neu5Gc MFI values obtained from three independent experiments. Within each experiment, MFI values were normalised to those of the corresponding WT control (WT = 1). Each point represents the mean of three technical replicate wells from one independent experiment. Data are presented as the mean ± SD (n = 3 independent experiments). CMAH(+) values were compared with the theoretical WT value of 1 using a two-tailed one-sample t-test, whereas paired untreated and mucinase-treated CMAH(+) values were compared using a two-tailed paired t-test. Holm’s correction was applied across the two comparisons. **:P < 0.01.

Exogenous Neu5Gc can be taken up by cells through pinocytosis and incorporated into cell-surface glycans through the sialic acid salvage pathway^22^. Furthermore, bacterial neuraminidase treatment reduced cell-surface Neu5Gc levels (Fig. 6), prompting us to investigate whether Neu5Gc produced following *CMAH* transfection could subsequently be acquired by neighbouring non-transfected cells. Considering mammalian neuraminidases such as cell surface NEU3 are upregulated in prostate cancers and implicated in regulating androgen signalling in LNCaP cells^59^, such a mechanism could constitute a “bystander effect”^60^ approach to realising sialome re-engineering in a prostate tumour microenvironment, without the requirement for transfection of all cells. Therefore to explore any potential “Neu5Gc-bystander effect”, a co-culture model experiment was performed using a mixture of CMAH (+) and non-transfected LNCaP cells to assess whether Neu5Gc incorporated into the glycans of transfected cells could subsequently result in increased Neu5Gc presentation on non-transfected adjacent cells. Non-transfected WT LNCaP cells were pre-labelled with a Far Red stain before co-culture with CMAH (+) LNCaP cells allowing both groups of cells to be distinguished and tracked over time using flow cytometry (Fig. 10a).

**Figure 10.**
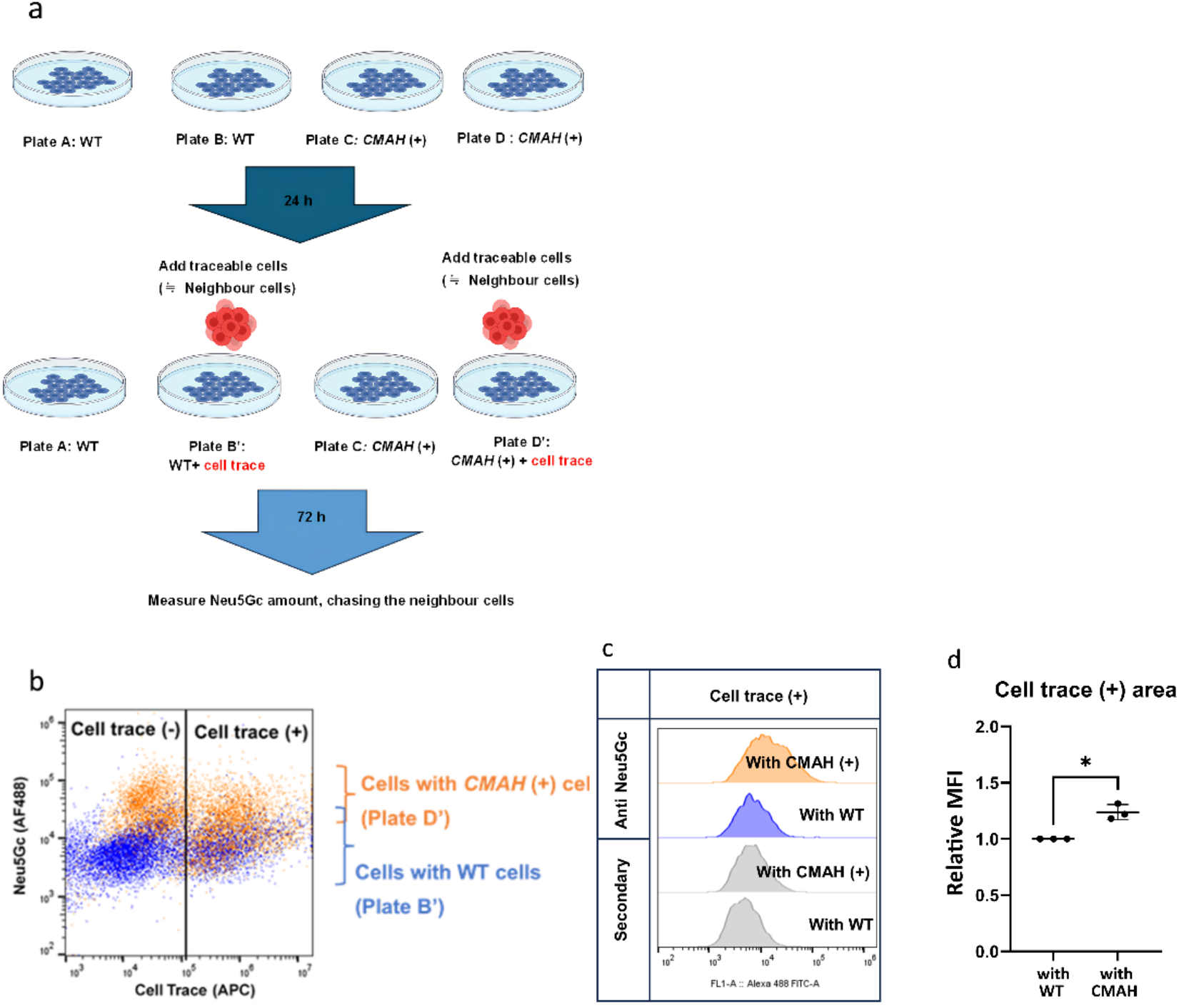
Cell trace analysis of Neu5Gc presentation in WT and CMAH (+) co-cultures. (a) Experimental workflow for assessing Neu5Gc presentation on WT, CMAH (+) cells, traced recipient LNCaP cells following co-culture with WT or CMAH (+) cells. WT cells were seeded in Plates A and B, whereas CMAH (+) cells were seeded in Plates C and D. After 24 h, cell traced recipient cells were added to Plates B and D, generating Plates Bʹ and Dʹ, respectively; Plates Aʹ and Cʹ were maintained without recipient cells as controls. After a further 72 h, all cells were harvested, and cell-surface Neu5Gc presentation was analysed by flow cytometry. (b) Overlay of representative flow-cytometry dot plots showing cell trace fluorescence in the APC channel and anti-Neu5Gc-associated AF488 fluorescence. The right-hand gate identifies the cell trace (+) recipient population. cell trace (+) recipient cells co-cultured with CMAH (+) cells (Plate Dʹ; orange) showed increased anti-Neu5Gc fluorescence compared with those co-cultured with WT cells (Plate Bʹ; blue). A representative overlay from three independent experiments is shown. (c) Representative flow-cytometry histograms showing anti-Neu5Gc-associated AF488 fluorescence within the gated cell trace (+) recipient population. Anti-Neu5Gc staining is shown for recipient cells co-cultured with CMAH (+) cells or WT cells, together with the corresponding secondary-antibody-only controls. (d) The graph presents anti-Neu5Gc MFI values for cell trace (+) recipient cells obtained from three independent experiments. Within each experiment, MFI values were normalised to those of recipient cells co-cultured with WT cells (WT co-culture = 1). Each point represents the mean of three technical replicate wells from one independent experiment. Data are presented as the mean ± SD (n = 3 independent experiments). Normalised values from recipient cells co-cultured with CMAH (+) cells were compared with the theoretical WT control value of 1 using a two-tailed one-sample *t*-test. *:*P* < 0.05.

Combined with the detection of the AF488 fluorescence signal from Neu5Gc staining, it was shown that WT cells did, in fact, incorporate some Neu5Gc on their surface. Briefly, cells were seeded as depicted in Fig. 10, WT in Plates A and B, and CMAH (+) cells in Plates C and D. On Day 3, additional WT cells, pre-stained with Cell trace Far Red (depicted as red cells in Fig. 10a), were added to Plates B and D, resulting in Plates Bʹ and Dʹ, respectively. After harvesting all cells, Neu5Gc cell surface presentation across Plates A, B’, C, and Dʹ was compared using the previously described anti-Neu5Gc flow cytometry assay. The resulting density plot overlays (Fig. 10b) illustrate Neu5Gc levels with and without cell trace staining. Notably CMAH (+) cells (left hand side of Fig. 10b, orange dots) show an upward shift compared to WT cells (blue dots), consistent with the expectation that CMAH (+) cells present higher levels of surface Neu5Gc. Whilst non-transfected cells stained with Cell trace Far Red (right side of Fig. 10b) added at day 3 to the CMAH (+) cell population clearly exhibited a higher level of Neu5Gc presentation than those added to WT cells (P < 0.05, Fig. 10c-d). Thus, these results confirm that Neu5Gc introduced through transfection can propagate in co-culture to neighbouring non-transfected cells, constituting a “Neu5Gc-bystander effect”.

## Discussion

In this study, transfection of LNCaP cells with the rat *CMAH* gene enabled the generation of a CMAH-expressing cell model for investigating Neu5Gc biosynthesis and cell-surface presentation. AlphaFold modelling predicted highly similar three-dimensional structures for rat and zebrafish CMAH, suggesting that the sequence differences between the two homologues did not substantially alter the overall protein fold. A cavity was identified close to the residues originally proposed to form the CMP-Neu5Ac-binding site. However, AlphaFold modelling alone cannot determine the precise substrate-binding mode, and structural analysis of CMAH in complex with CMP-Neu5Ac would be required to define these interactions. Consistent with the similarity of the predicted structures, flow-cytometry analysis showed comparable levels of cell-surface Neu5Gc following transfection with the rat and zebrafish *CMAH* plasmids. Future studies combining molecular docking with experimental validation could provide further insight into substrate recognition and enzyme function.

Time-course analysis of LNCaP *CMAH* transfection indicated that rat *CMAH* transfection effectively altered cell-surface sialic acid composition. Notably, the 96-hour time point showed the greatest shift, suggesting that Neu5Gc expression on the cell surface was highest among the time points examined. By 120 hours, Neu5Gc levels appeared to decrease, possibly due to shedding from the cell surface or because transgene expression or Neu5Gc production had reached a plateau. Although rat *CMAH* transfection clearly increased cell-surface Neu5Gc levels, the detailed metabolic fate of Neu5Gc remains unclear. In a related experiment conducted by Bergfeld *et al.*, human THP-1 cells were pulsed with exogenous Neu5Gc for three days and subsequently cultured under Neu5Gc-free conditions to observe the fate of the incorporated Neu5Gc over time^61^. DMB-HPLC analysis revealed that approximately 55% of total sialic acids were Neu5Gc after the initial pulse. During the chase period, Neu5Gc levels declined. By day five, about half of the incorporated Neu5Gc had been lost, which is broadly consistent with the reduction observed in the present study. Bergfeld *et al.* also proposed that evolutionary pressures tend to favour energy-efficient systems. Accordingly, cells capable of synthesising specific molecules may also possess the enzymatic machinery required for their degradation and recycling. Following sialidase treatment, Neu5Gc expression on CMAH (+) LNCaP cells was significantly reduced, indicating that Neu5Gc was presented as terminal cell-surface sialic acids accessible to enzymatic cleavage. Combined with the inhibitory effect of benzyl-α-GalNAc, these findings suggest that Neu5Gc is incorporated predominantly into O-glycans. Supporting this, Valenzuela *et al*. reported that benzyl-α-GalNAc treatment in LNCaP cells led to loss of O-glycan sialylation^45^. Overall, the findings suggest that Neu5Gc incorporation predominantly occurs on O-glycans in transfected LNCaP cells. This observation is particularly relevant because aberrantly glycosylated tumour-associated mucins, which are rich in O-glycans, have been implicated in promoting immune suppression within the tumour microenvironment. Therefore, the preferential incorporation of Neu5Gc into O-glycans may have important implications for tumour–immune interactions and warrants further investigation.

For more definitive characterisation, glycopeptide analysis of WT and CMAH (+) cells was conducted. The glycan profile of WT LNCaP cells revealed several well-characterised glycopeptides bearing Neu5Ac-containing glycans (Supplementary Table S1). A comparative analysis by Shah *et al*. (2015) used integrated global proteomics and glycoproteomics to examine PC3 and LNCaP cells. Their study identified and quantified N-linked glycopeptides bearing glycans in iTRAQ (isobaric tags for relative and absolute quantification)-labelled samples. This analysis revealed 67 distinct glycan compositions. However, Shah *et al*. did not detect any sialylated N-glycans. Similarly, in our dataset, only the N-glycan structure HexNAc(4)Hex(4)Fuc(2)Neu5Ac(1) was identified in WT cells. This limited detection of sialylated N-glycans may reflect methodological differences, biological variation, or the low abundance of these structures in LNCaP cells. Although total cellular glycopeptides were successfully identified, flow cytometry specifically detects sialylated glycoconjugates exposed on the cell surface. Accordingly, only one cell-surface glycopeptide, transferrin receptor protein 1 (TFRC), carrying a Neu5Ac-containing glycan, was identified in the WT cell dataset.

Nevertheless, comparison of the WT and CMAH (+) glycoproteomic datasets indicated a partial replacement of Neu5Ac by Neu5Gc on several glycopeptides. To further evaluate the coverage of cell-surface proteins, our global proteomic dataset was compared with the dataset reported by Shah *et al*. using STRING. In their dataset, which contained the 100 most abundant proteins, 11 proteins, including FOLH1, EFNA5, CALR, CD47, APMAP, SORT1, ALCAM, LAMP1, IGF2R, PLXNB2, and TFRC, were predicted to localise to the plasma membrane, nine of which were reported to be sialylated. The limited detection of these proteins in the present study suggests incomplete coverage of the cell-surface glycoproteome and indicates that some sialylated cell-surface glycopeptides may have been lost during sample preparation or were present below the detection limit of the analytical workflow.

This interpretation is supported by the flow cytometry results, which showed that mucinase treatment reduced Neu5Gc expression on CMAH (+) cells, suggesting that mucin-type glycoproteins are the predominant carriers of Neu5Gc. Consequently, the current glycoproteomic workflow may require further optimisation to improve the recovery of cell-surface glycoproteins. Future studies could incorporate mucinase digestion prior to glycopeptide analysis to improve the characterisation of mucin-associated glycans. In addition, membrane enrichment techniques may minimise the loss of plasma membrane proteins during sample preparation. For example, the membrane fractionation protocol described by Sadler *et al*. has been shown to enrich total membrane proteins and may improve the identification of cell-surface glycoproteins^62^. Similarly, a ball-bearing homogeniser, as described by Thomas *et al*., could be used to gently homogenise LNCaP cells before glycopeptide analysis^63^. This method has been shown to enhance protein extraction while preserving membrane-associated proteins, thereby improving downstream glycoproteomic analysis. Together, these methodological refinements could increase the recovery and detection of cell-surface glycoproteins, enabling a more comprehensive characterisation of Neu5Gc-containing glycoconjugates.

Finally, co-culture experiments demonstrated that Neu5Gc generated following *CMAH* transfection could be transferred to neighbouring prostate cancer cells and subsequently incorporated into their cell-surface glycans. These findings highlight not only the effects of Neu5Gc expression in transfected cells but also its potential influence on the surrounding tumour microenvironment. Based on these observations, we propose that Neu5Gc-containing molecules may be released from donor cells, potentially through the activity of endogenous sialidases, some of which are dysregulated in prostate and other cancers^64^. In this scenario the liberated Neu5Gc may be taken up by neighbouring tumour cells, activated through the sialic acid salvage pathway, and reincorporated into cell-surface glycans, establishing a self-propagating mechanism of Neu5Gc dissemination within the tumour microenvironment (Fig. 11). This may represent a putative Neu5Gc bystander effect, whereby non-transfected cells acquire a Neu5Gc-positive phenotype through intercellular transfer rather than direct genetic modification. However, further experiments are required to identify the molecular form in which Neu5Gc is transferred and to confirm the involvement of sialidases and the sialic acid salvage pathway.

**Figure 11.**
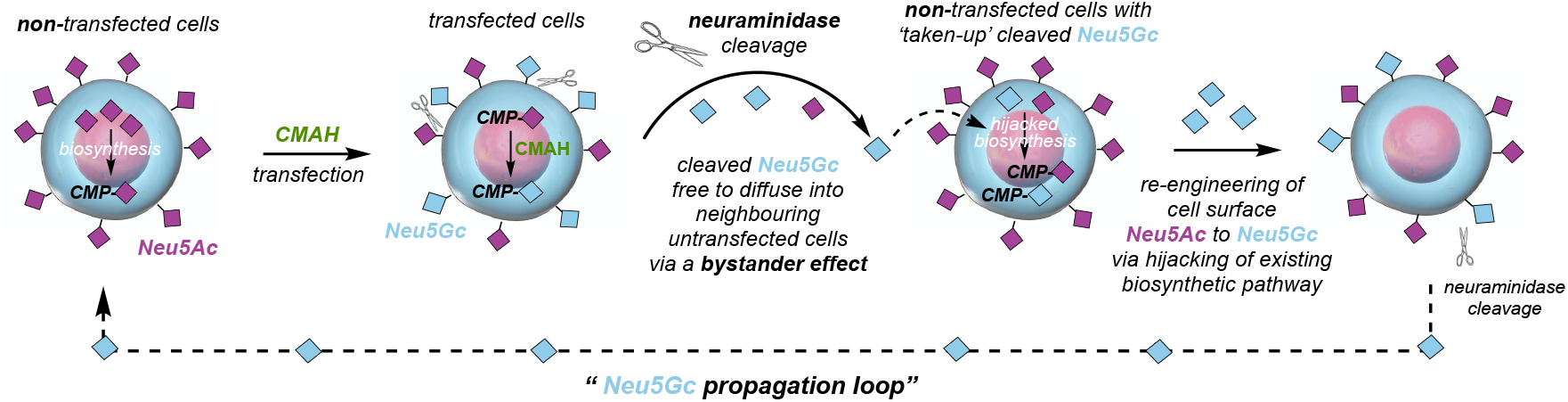
Schematic representation of the proposed propagation of Neu5Gc through a “bystander effect”. Following CMAH transfection, the cell surface becomes decorated with a mixture of Neu5Ac and Neu5Gc. Neuraminidases such as NEU3 cleave both sialic acids from the cell surface, allowing Neu5Gc to diffuse into neighbouring non-transfected cells via a “bystander effect”. Neu5Gc can then enter the native sialic acid biosynthetic pathway in recipient cells, leading to intracellular CMP-Neu5Gc formation. Consequently, both CMAH (+) and neighbouring non-transfected cells undergo progressive remodelling of cell surface sialylation, forming a potential Neu5Gc propagation loop.

A related form of microenvironment-mediated glycan regulation was reported by Egan and colleagues.^65^ In that study, murine mesenchymal stromal cells (MSCs) were conditioned with the secretome of CT26 tumour cells (CT26-MSCs^TCS^). Flow cytometric analysis demonstrated significantly increased Sambucus nigra agglutinin I SNA-I binding in the conditioned MSCs, indicating an increase in α2,6-linked sialylation. Thus, this study showed that tumour-derived soluble factors can modify the sialylation phenotype of neighbouring stromal cells. Similarly, the Neu5Gc transfer observed in the present study suggests that interactions between neighbouring tumour cells may influence their glycosylation patterns.

Tumours are composed of genetically and phenotypically heterogeneous cell populations. A Neu5Gc bystander mechanism could therefore extend a Neu5Gc-positive glycosylation phenotype to cells that have not themselves undergone *CMAH* transfection, potentially reducing intratumoral glycosylation heterogeneity independently of genetic modification. This concept is relevant to tumour glycobiology because it suggests that glycosylation is determined not only by the intrinsic biosynthetic machinery of individual tumour cells but also by intercellular interactions within the tumour microenvironment.

These findings also have important implications for studies of sialic acid–immune interactions. Humans lack functional CMAH and cannot synthesise Neu5Gc endogenously, whereas wild-type mice express Neu5Gc naturally. Moreover, Neu5Gc present in animal-derived serum can be taken up and incorporated into the glycans of cultured human cells. Consequently, the use of human serum in cell-culture experiments and, where appropriate, CMAH-deficient mouse models may help distinguish experimentally induced or transferred Neu5Gc from Neu5Gc originating from serum or endogenous murine CMAH activity.

Our experimental model therefore provides a platform for investigating how Neu5Ac-to-Neu5Gc sialome re-engineering influences prostate cancer cell biology. It also offers a system for examining the mechanisms responsible for Neu5Gc transfer between tumour cells and the potential consequences of this transfer for tumour progression. Given the established role of aberrant cell-surface sialylation in tumour immune evasion, this model could facilitate future studies of how Neu5Gc affects tumour–immune interactions. Such studies may determine whether the replacement of Neu5Ac with Neu5Gc alters immune recognition and contributes to the establishment or modulation of an immunosuppressive tumour microenvironment.

## Methods

All materials and reagents were purchased from Thermo Fisher Scientific or Sigma, and chemicals were sourced from Merck, unless otherwise stated.

### cDNA clones and plasmid stock preparation

Rat and zebrafish *CMAH* cDNA clones (ABIN4054902, ABIN4057525) in the pExpress1 vector were purchased from Abnova (Limerick, USA). Less than 1 µL of glycerol stock containing the cDNA clone was streaked onto 20 mL LB agar containing 100 μg/mL ampicillin, followed by incubation at 37 °C overnight. A 4 mL starter culture grown from a single colony was transferred into 400 mL LB medium containing 100 μg/mL ampicillin and incubated at 37 °C overnight with shaking at 200 rpm. The culture was harvested by centrifugation (4,000 × g, 4 °C, 30 min), and plasmid DNA was extracted using the EndoFree® Plasmid Purification Maxi Kit (QIAGEN, Hilden, Germany) according to the manufacturer’s protocol. DNA was eluted in 500 µL of the supplied Buffer TE.

### CMAH sequences comparison

Multiple sequence alignment of CMAH proteins was performed using the GENETYX parallel editor (GENETYX Corporation, Tokyo, Japan), incorporating amino acid sequences from chimpanzee (NP_001009041), human (AAC68881), pig (NP_001106486), mouse (NP_001104580), rat (NP_001019444.1), and zebrafish (NP_001002192).

### Three-dimensional structural analysis of rat and zebrafish CMAH proteins

The protein structure prediction AI system AlphaFold2, developed by DeepMind (Google DeepMind), was accessed through the Google Colab notebook interface. Protein structures were predicted using ColabFold. The resulting Protein Data Bank (PDB) files were visualised and analysed using PyMOL (LCC, Schrödinger, Germany), a molecular visualisation tool based on Python. Structures were visualised as molecular surfaces, cartoon representations of secondary structure, and atomic models.

### Cell culture

The androgen-sensitive prostate cancer cell line LNCaP was routinely maintained in Gibco RPMI-1640 medium in 75 cm² flasks. RPMI-1640 was supplemented with 10% heat-inactivated fetal calf serum (FCS), 1% penicillin–streptomycin, and 1% L-glutamine.

Cells were incubated at 37 °C in a humidified atmosphere containing 5% CO₂. For routine passaging or experimental procedures, cells were detached using either Trypsin/EDTA (Trypsin) or 10 mM ethylenediaminetetraacetic acid (EDTA) prepared in phosphate-buffered saline (PBS), as appropriate.

### *CMAH* gene transfection

Cells were seeded in 6-well plates at a density of 200,000 cells per well in 2 mL of standard culture medium containing FCS. On Day 2, transfection was performed using either rat or zebrafish CMAH plasmids with X-tremeGENE HP DNA Transfection Reagent (Roche, Penzberg, Germany), following the manufacturer’s protocol. A 3:1 (v/w) ratio of transfection reagent to plasmid DNA was used (2 µg DNA and 6 µL reagent per well). On Day 3, the culture medium was replaced with fresh medium containing 5% human serum (type AB) to prevent further incorporation of exogenous Neu5Gc. Cells were then incubated for a total of 96 hours post-transfection. On Day 6, cells were detached using EDTA and processed for flow cytometry to assess Neu5Gc expression.

### Flow cytometry

Harvested cells were washed with PBS by centrifugation (1,500 × g for 5 min at 4 °C) and seeded into 96-well plates at approximately 70,000 cells per well. Cells were maintained at 4 °C throughout the washing, staining, and sample-preparation steps. The plated cells were gently washed with PBS and incubated for 1 h with purified chicken anti-Neu5Gc primary antibody (BioLegend, CA, USA; 1:500 in PBS). Following three washes with PBS, cells were incubated for 1 h with goat anti-chicken IgY Alexa Fluor 488-conjugated secondary antibody (Invitrogen, CA, USA; 1:200 in PBS). After three further washes with PBS, cells were fixed overnight in PBS containing 1% formaldehyde and subsequently resuspended in PBS for flow cytometry analysis. Fluorescence was measured using a CytoFLEX S flow cytometer (Beckman Coulter, CA, USA), with at least 5,000 events acquired per sample. Neu5Gc expression was quantified as the mean fluorescence intensity (MFI) of Alexa Fluor 488 within the gated cell population. Each independent experiment included three technical replicate wells. Unless otherwise stated in the figure legends, the MFI values from the three wells were averaged to obtain a single value representing one independent experiment. Data are presented as the mean ± standard deviation (SD) from three independent experiments (*n* = 3). To enable comparisons across independent experiments, the MFI of each sample was normalised to that of the corresponding control condition within the same experiment. For experiments involving three or more matched conditions, a repeated-measures one-way ANOVA followed by the multiple-comparisons test specified in the corresponding figure legend was used. Statistical significance was defined as *P* < 0.05.

### Chemical and enzymatic treatments for Neu5Gc measurement

#### O-and N-linked glycosylation inhibitor treatment

LNCaP cells were transfected with the rat *CMAH* gene on Day 2 after cell seeding, and R10 medium (RPMI-1640 + 10% FBS) was replaced with R5-HS (RPMI-1640 + 5% human serum) on Day 3. Immediately after the medium change, cells were treated with either 2 mM of the O-linked glycosylation inhibitor Benzyl 2-acetamido-2-deoxy-α-D-galactopyranoside (Benzyl-α-GalNAc) or 25 μM of the N-linked glycosylation inhibitor deoxymannojirimycin (dMNJ), dissolved in PBS. Cells were incubated at 37 °C for 72 hours in the presence of these inhibitors. On Day 6, cells were harvested and prepared for flow cytometry analysis to quantify Neu5Gc expression.

#### Mucinase and sialidase treatment

LNCaP cells were transfected with the rat *CMAH* gene, and the medium was changed as described above. On Day 6, following harvest, cells were resuspended in PBS and incubated with the appropriate enzymatic reagents: IMPa at a concentration of 1 U per 10 µg of glycoprotein, and StcE at 10 µg per 1 × 10⁶ cells for 5 hours at 37 °C, or neuraminidase (1 U per 10 mg/ml incubation mixture) for 1 h at 37 °C. After enzymatic incubation, cells were processed for flow cytometry analysis to assess Neu5Gc expression levels.

### Cell trace analysis

Cell staining was conducted at room temperature. Harvested LNCaP cells were washed by centrifugation with R10. To remove serum, cells were resuspended in R0 (RPMI-1640 without serum) at 1 × 10⁶ cells/mL and stained with 2 µMcell trace™ Far Red DDAO-SE (1:1000 dilution), mixed by inversion, and protected from light. During the 20 min incubation at room temperature, the sample was inverted regularly. To remove free dye, five times the original volume of R10 was added and incubated for 5 min, followed by three washing steps with fresh R5-HS. Cell trace-stained LNCaP cells were added to non-transfected or rat *CMAH (+)* LNCaP cells 24 hours after transfection. At the same time, the medium was changed from R10 to R5-HS. On Day 6, stained and unstained cells were processed for Neu5Gc staining and analysed using a CytoFLEX S, as described above, with FITC and APC MFIs recorded.

### Glycopeptide analysis

For sample preparation, approximately 1.0 × 10⁶ cells (50–100 µg glycoprotein) were harvested using PBS and gentle pipetting. The cell pellet was lysed in five volumes (v/v) of freshly prepared sodium dodecyl sulfate (SDS) buffer at room temperature. The mixture was pipetted up and down several times to aid lysis. Lysates were not cooled on ice, as this can cause precipitation. Glycopeptide analysis was performed by Dr Adam Dowle at the Metabolomics & Proteomics Laboratory, Bioscience Technology Facility, University of York. Protein lysates were digested using the S-Trap protocol following reduction with tris(2-carboxyethyl)phosphine (TCEP) and alkylation with methyl methanethiosulfonate. A combination of trypsin and Lys-C proteases was used for digestion. The resulting peptides were analysed by liquid chromatography–mass spectrometry (LC-MS) over a 2 h acquisition period. Peptide separation was performed using a 50 cm EASY-Spray PepMap™ column with flow delivered by an mClass UPLC system (Waters, Milford, USA). Data were acquired on an Orbitrap Fusion Tribrid mass spectrometer operated in data-dependent acquisition (DDA) mode. High-resolution MS1 spectra were acquired in the Orbitrap analyser, and rapid MS2 spectra were collected following higher-energy collisional dissociation (HCD) in the ion trap analyser using a 1 s top-speed cycle. Spectra were searched against the human subset of the SwissProt database using the Byonic search engine^66^. β-Methylthiolation was set as a fixed modification to account for alkylation, while oxidation and N-terminal acetylation were included as common variable modifications. Protein-level results were filtered to include only those with at least two unique peptides and a global false discovery rate (FDR) of 1%, determined empirically using a decoy database. Peptide-level data were similarly filtered at 1% FDR and restricted to individual peptide-spectrum matches with a log probability score greater than 1.3 (P < 0.05).

## Data availability statement

The datasets generated and/or analysed during the current study are included in this published article and its Supplementary Information files. Additional data are available from the corresponding author upon reasonable request.

## Acknowledgements

We thank Dr. Adam Dowle and The York Centre of Excellence in Mass Spectrometry. The York Centre of Excellence in Mass Spectrometry was created thanks to a major capital investment through Science City York, supported by Yorkshire Forward with funds from the Northern Way Initiative, and subsequent support from EPSRC (EP/K039660/1; EP/M028127/1). This work was supported by the University of York Department of Chemistry, a Horizon Europe Guarantee Consolidator award to M.A.F. (selected by the ERC, funded by UKRI; EP/X023680/1), and an MRC DiMeN Doctoral Training Partnership PhD award to E.H.

## Author Contributions

M.A.F. and N.S conceived the study. Y.U., A.R.N. and E.H. performed the experiments. M.A.F. and N.S. contributed to study design and data interpretation. Y.U., N.S. and M.A.F. drafted the manuscript. All authors reviewed and approved the final manuscript.

