## Supplementary information for "Characterisation of prostate cancer sialome re-engineering via *CMAH* transfection reveals a bystander effect that propagates Neu5Gc presentation to neighbouring cells"

### Supplementary Figures S1–S4

Quality assessment of the Byonic database search for glycoproteomic analysis of WT and *CMAH*(+) LNCaP cells. Protein score plots show forward (target) and reverse (decoy) protein identifications used to assess the false discovery rate (FDR). Precursor mass error plots show the difference between observed and calculated precursor masses for all forward spectrum identifications. The distribution of precursor mass errors around zero indicates good mass accuracy and calibration of the LC–MS/MS data.

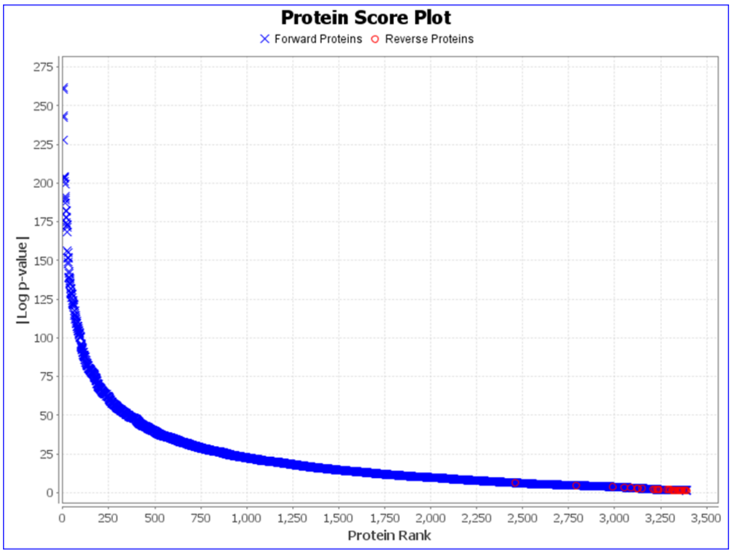


**Supplementary Figure S1.**

Byonic protein score plot of WT LNCaP cells. Forward protein identifications (blue crosses) and reverse (decoy) protein identifications (red circles) are plotted according to protein rank. The y-axis represents the absolute base-10 logarithm of the P-value (|Log P-value|), corresponding to the Log Prob values reported in Supplementary Tables S1 and S2. The clear separation between target and decoy matches indicates a low false discovery rate (FDR) and high confidence in protein identifications.

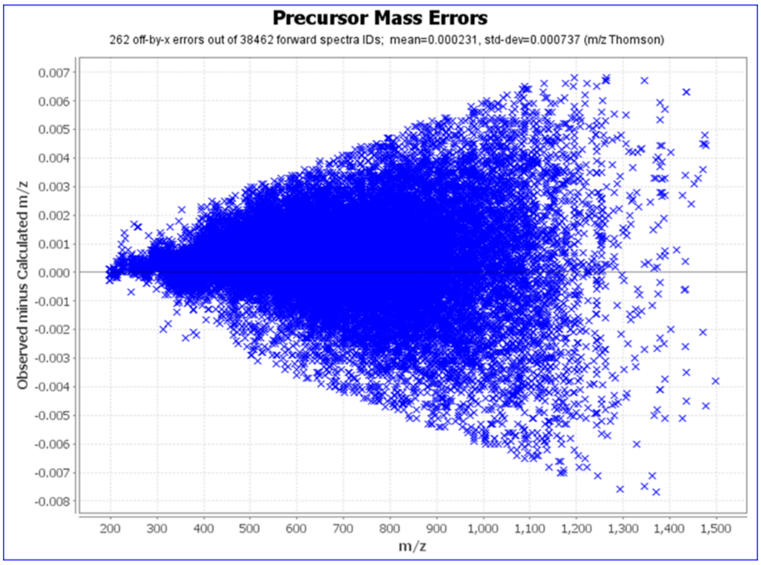


**Supplementary Figure S2.**

Precursor mass-error distribution for WT LNCaP peptide spectra. The precursor mass error, calculated as the observed m/z minus the calculated m/z, is plotted against precursor m/z for all forward spectrum identifications generated by Byonic. A total of 38,462 forward spectra were identified, including 262 off-by-x mass-assignment errors. The mean precursor mass error was 0.000231 Th, with a standard deviation of 0.000737 Th.

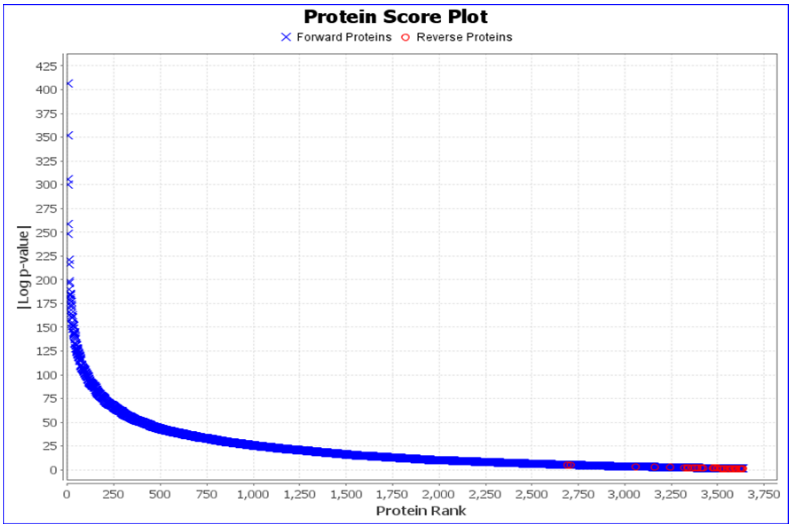


**Supplementary Figure S3.**

Byonic protein score plot of CMAH(+) LNCaP cells. Forward protein identifications (blue crosses) and reverse (decoy) protein identifications (red circles) are plotted according to protein rank. The y-axis represents the absolute base-10 logarithm of the P-value (|Log P-value|), corresponding to the Log Prob values reported in Supplementary Tables S1 and S2.

The clear separation between target and decoy matches indicates a low false discovery rate (FDR) and high confidence in protein identifications.

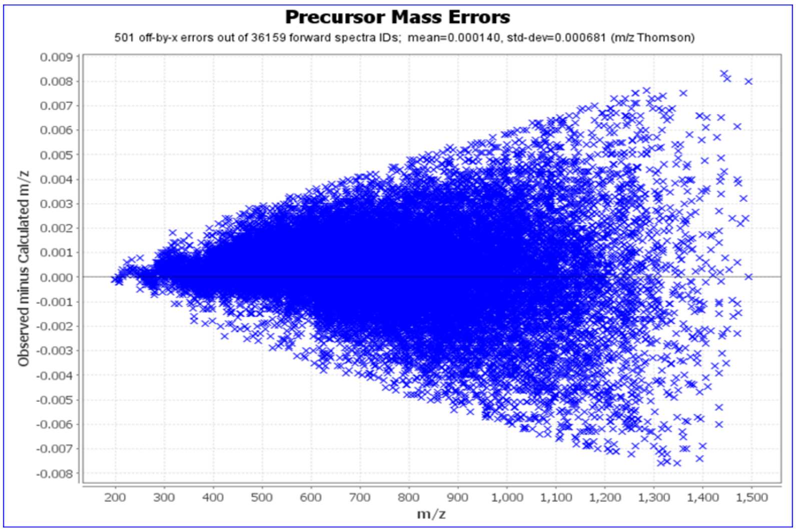


**Supplementary Figure S4.**

Precursor mass-error distribution for CMAH(+) LNCaP peptide spectra. The precursor mass error, calculated as the observed m/z minus the calculated m/z, is plotted against precursor m/z for all forward spectrum identifications generated by Byonic. A total of 36,159 forward spectra were identified, including 501 off-by-x mass-assignment errors. The mean precursor mass error was −0.000140 Th, with a standard deviation of 0.000681 Th.

**Supplementary Figures S5–S13**

Representative MS/MS spectra of glycopeptides identified in wild-type (WT) and CMAH(+) LNCaP cells. Glycopeptides were identified using Byonic and confirmed by manual inspection of the MS/MS spectra. Fragment ion assignments are shown in blue (b ions), red (y ions), and green (glycan-related ions, where applicable).

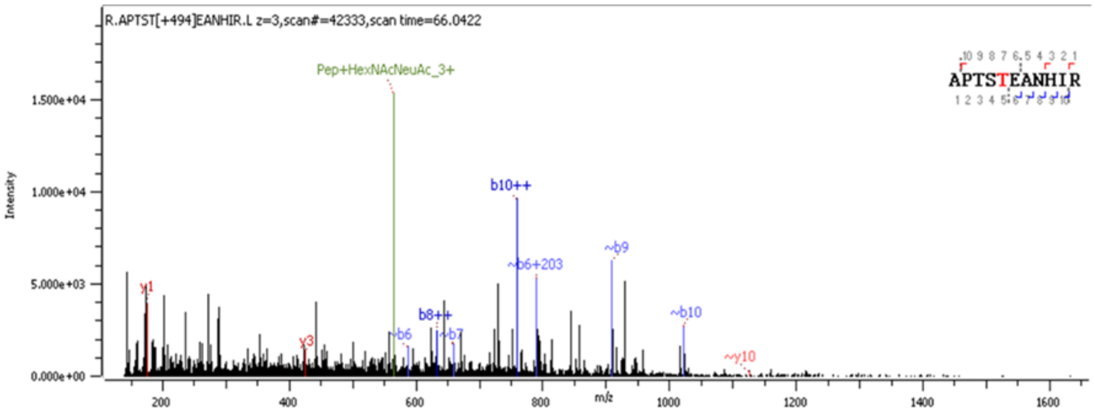


**Supplementary Figure S5.**

Representative MS/MS spectrum of the WT LNCaP glycopeptide R.APTST[+494.175]EANHIR.L carrying HexNAc(1)NeuAc(1). The precursor ion (z = 3, scan 42333, scan time = 66.0422 min) corresponds to the tryptic peptide APTSTEANHIR modified with a HexNAc(1)NeuAc(1) glycan attached to Thr5.

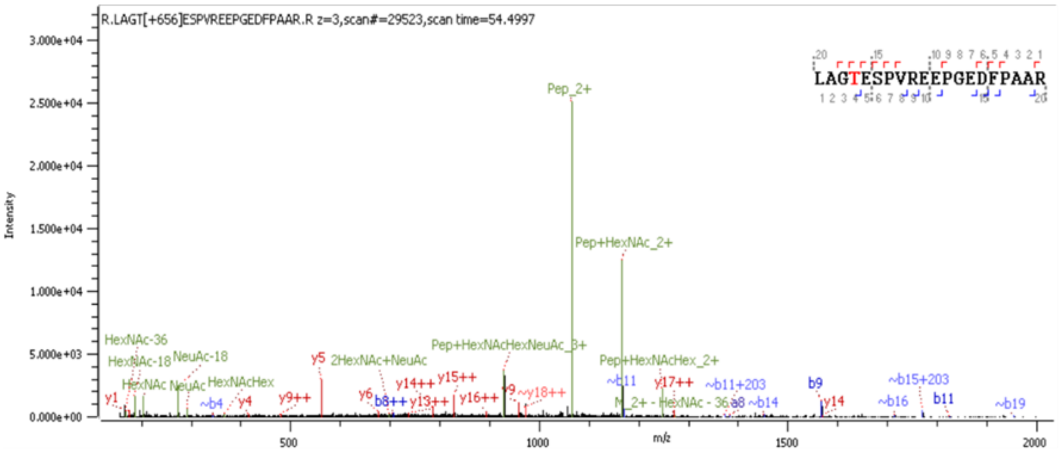


**Supplementary Figure S6.**

Representative MS/MS spectrum of the WT LNCaP glycopeptide R.LAGT[+656.228]ESPVREEPGEDFPAAR.R carrying HexNAc(1)Hex(1)NeuAc(1). The precursor ion (z = 3, scan 29523, scan time = 54.4997 min) corresponds to the tryptic peptide LAGTESPVREEPGEDFPAAR modified with a HexNAc(1)Hex(1)NeuAc(1) glycan attached to Thr4.

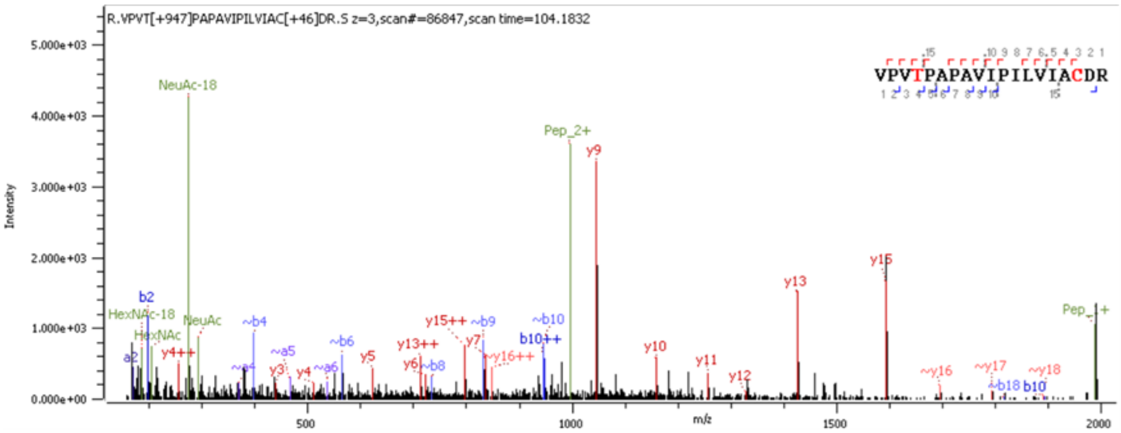


**Supplementary Figure S7.**

Representative MS/MS spectrum of the WT LNCaP glycopeptide R.VPVT[+947.323]PAPAVIPILVIAC[+45.988]DR.S carrying HexNAc(1)Hex(1)NeuAc(2). The precursor ion (z = 3, scan 86847, scan time = 104.1832 min) corresponds to the tryptic peptide VPVTPAPAVIPILVIACDR modified with a HexNAc(1)Hex(1)NeuAc(2) glycan attached to Thr4. The cysteine residue was β-methylthiolated (+45.988 Da) as a fixed modification.

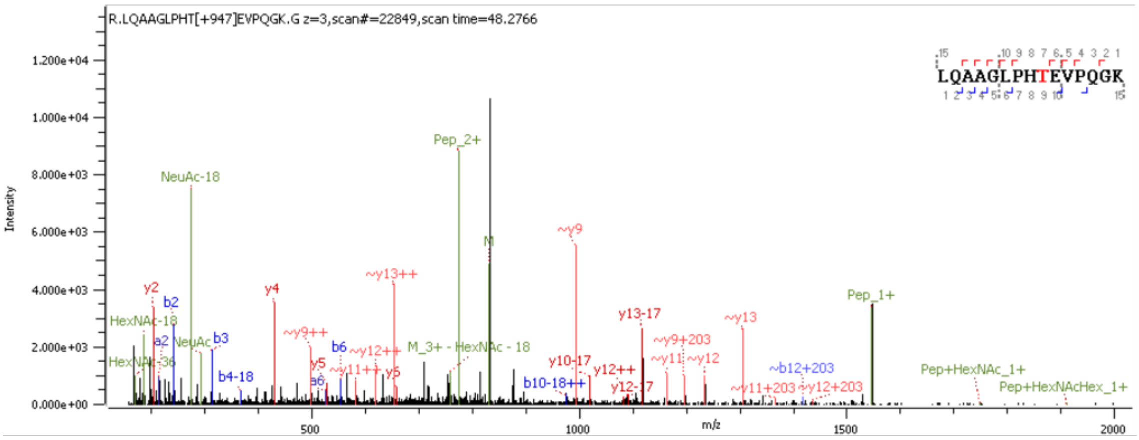


**Supplementary Figure S8.**

Representative MS/MS spectrum of the WT LNCaP glycopeptide R.LQAAGLPHT[+947.323]EVPQGK.G carrying HexNAc(1)Hex(1)NeuAc(2). The precursor ion (z = 3, scan 22849, scan time = 48.2766 min) corresponds to the tryptic peptide LQAAGLPHTEVPQGK modified with a HexNAc(1)Hex(1)NeuAc(2) glycan attached to Thr9.

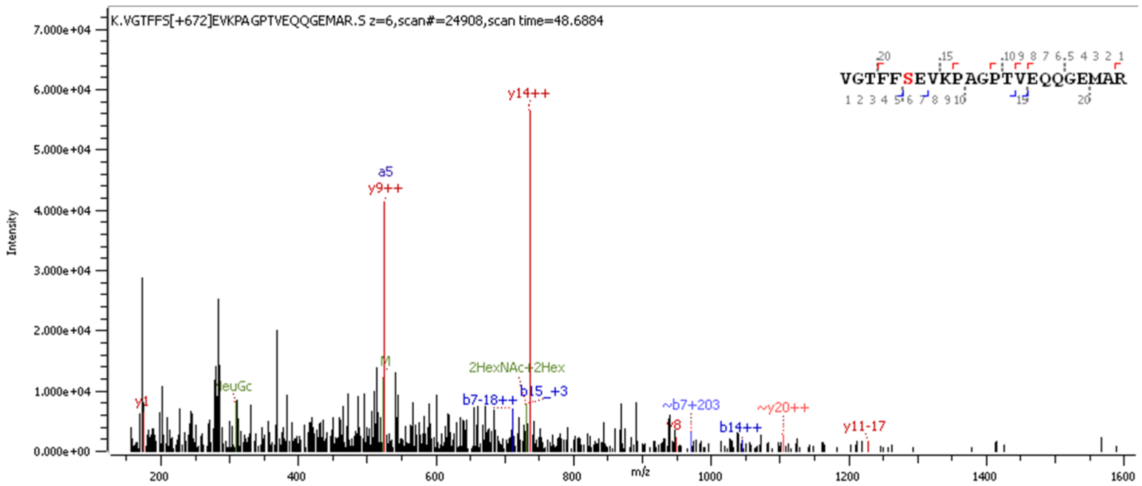


**Supplementary Figure S9.**

Representative MS/MS spectrum of the CMAH(+) LNCaP glycopeptide K.VGTFFS[+672.223]EVKPAGPTVEQQGEMAR.S carrying HexNAc(1)Hex(1)NeuGc(1). The precursor ion (z = 6, scan 24908, scan time = 48.6884 min) corresponds to the tryptic peptide VGTFFSEVKPAGPTVEQQGEMAR modified with a HexNAc(1)Hex(1)NeuGc(1) glycan attached to Ser6.

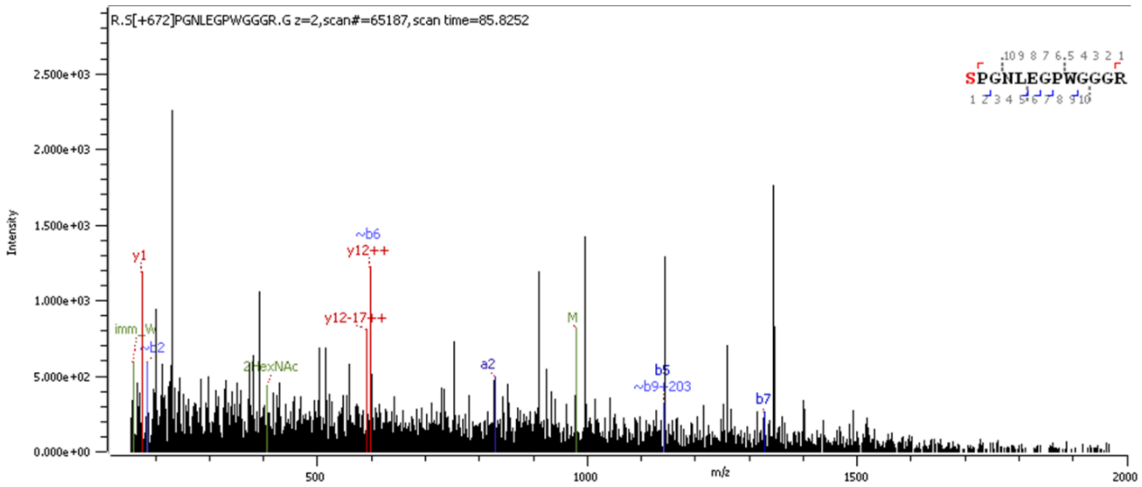


**Supplementary Figure S10.**

Representative MS/MS spectrum of the CMAH(+) LNCaP glycopeptide R.S[+672.223]PGNLEGPWGGGR.G carrying HexNAc(1)Hex(1)NeuGc(1). The precursor ion (z = 2, scan 65187, scan time = 85.8252 min) corresponds to the tryptic peptide SPGNLEGPWGGGR modified with a HexNAc(1)Hex(1)NeuGc(1) glycan attached to Ser1.

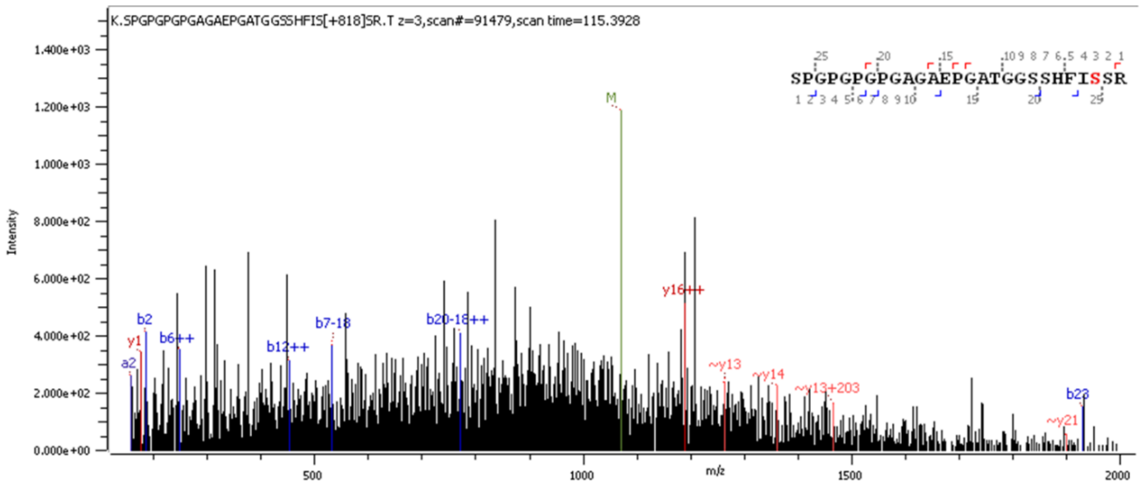


**Supplementary Figure S11.**

Representative MS/MS spectrum of the CMAH(+) LNCaP glycopeptide K.SPGPGPGPGAGAEPGATGGSSHFIS[+818.280]SR.T carrying HexNAc(1)Hex(1)Fuc(1)NeuGc(1). The precursor ion (z = 3, scan 91479, scan time = 115.3928 min) corresponds to the tryptic peptide SPGPGPGPGAGAEPGATGGSSHFISSR modified with a HexNAc(1)Hex(1)Fuc(1)NeuGc(1) glycan attached to Ser25.

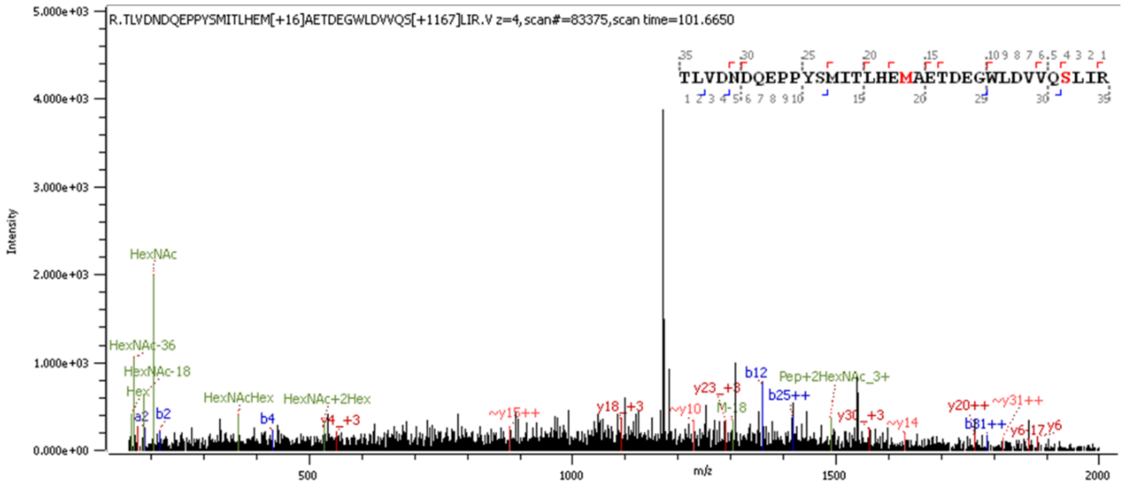


**Supplementary Figure S12.**

Representative MS/MS spectrum of the CMAH(+) LNCaP glycopeptide R.TLVDNDQEPPYSMITLHEM[+15.995]AETDEGWLDVVQS[+1167.418]LIR.V carrying HexNAc(2)Hex(1)Fuc(2)NeuGc(1). The precursor ion (z = 4, scan 83375, scan time = 101.6650 min) corresponds to the tryptic peptide TLVDNDQEPPYSMITLHEMAETDEGWLDVVQSLIR modified with a HexNAc(2)Hex(1)Fuc(2)NeuGc(1) glycan attached to Ser32. Oxidation of methionine (+15.995 Da) was included as a variable modification.

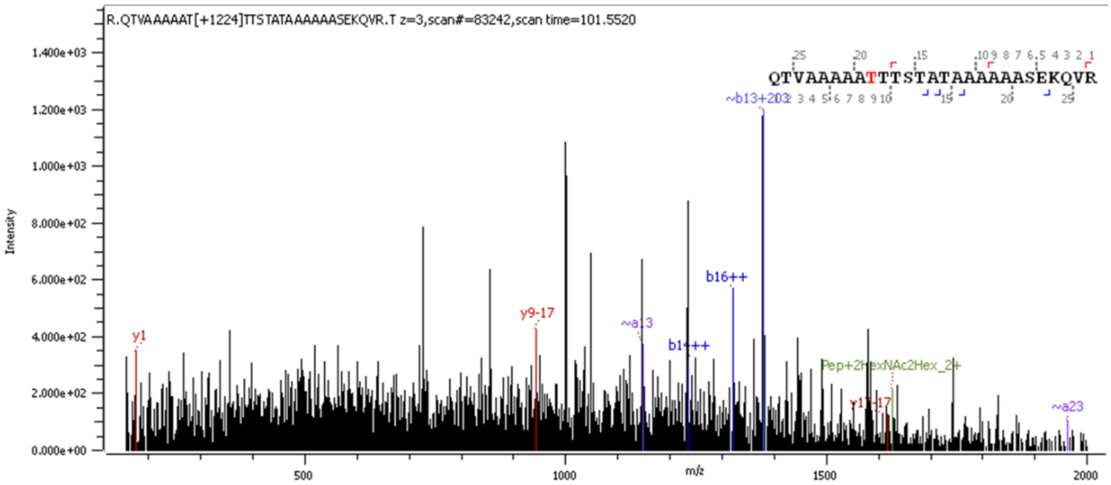


**Supplementary Figure S13.**

Representative MS/MS spectrum of the CMAH(+) LNCaP glycopeptide R.QTVAAAAAT[+1224.439]TTSTATAAAAAASEKQVR.T carrying HexNAc(3)Hex(1)Fuc(1)NeuGc(1). The precursor ion (z = 3, scan 83242, scan time = 101.5520 min) corresponds to the tryptic peptide QTVAAAAATTTSTATAAAAAASEKQVR modified with a HexNAc(3)Hex(1)Fuc(1)NeuGc(1) glycan attached to Thr9.

**Supplementary Table S1. Glycopeptides with sialic acids in WT LNCaP cells**

| UniProt | Protein Name | Glycans | O/N | Location | Log Prob |
|---|---|---|---|---|---|
| Q8NBJ4 | GOLM1_HUMAN Golgi membrane protein 1 | HexNAc(1)Hex(1)NeuAc(2) | O | Cytoplasm, endoplasmic reticulum | 14.28 |
| Q8NBJ4 | GOLM1_HUMAN Golgi membrane protein 1 | HexNAc(2)Hex(2)Fuc(2)NeuAc(1) | O | Cytoplasm, endoplasmic reticulum | 13.01 |
| P26572 | MGAT1_HUMAN Alpha-1,3-mannosyl-glycoprotein 2-beta-N-acetylglucosaminyltransferase | HexNAc(1)Hex(1)NeuAc(2) | O | Golgi apparatus membrane, Cytoplasm, perinuclear region | 12.22 |
| Q8NBJ4 | GOLM1_HUMAN Golgi membrane protein 1 | HexNAc(2)Hex(2)Fuc(1)NeuAc(1) | O | Cytoplasm, endoplasmic reticulum | 11.71 |
| P02786 | TFR1_HUMAN Transferrin receptor protein 1 | HexNAc(1)Hex(1)NeuAc(1) | O | Cell surface | 9.13 |
| Q8IYS2 | K2013_HUMAN Uncharacterised protein KIAA2013 | HexNAc(2)Hex(2)Fuc(2)NeuAc(1) | O | Membrane | 8.07 |
| Q9HC07 | TM165_HUMAN Transmembrane protein 165 | HexNAc(1)Hex(1)NeuAc(2) | O | Golgi apparatus membrane, golgi apparatus, trans-golgi network membrane | 6.72 |
| Q9BRK5 | CAB45_HUMAN 45 kDa calcium-binding protein | HexNAc(1)Hex(1)NeuAc(2) | O | Golgi apparatus lumen, cytoplasm, cell membrane | 6.35 |
| P02786 | TFR1_HUMAN Transferrin receptor protein 1 | HexNAc(1)Hex(1)NeuAc(2) | O | Cell surface | 5.47 |
| Q8IYS2 | K2013_HUMAN Uncharacterised protein KIAA2013 | HexNAc(1)Hex(1)NeuAc(2) | O | Membrane | 5.22 |
| O95947 | TBX6_HUMAN T-box transcription factor TBX6 | HexNAc(3)Hex(3)Fuc(1)NeuAc(2) | O | Nucleus, secreted | 5.15 |
| Q6SPF0 | SAMD1_HUMAN Atherin | HexNAc(2)Hex(2)Fuc(1)NeuAc(2) | O | Nucleus | 3.94 |
| O95674 | CDS2_HUMAN Phosphatidate cytidylyltransferase 2 | HexNAc(2)Hex(2)Fuc(2)NeuAc(1) | O | Nucleus | 3.48 |
| P18124 | RL7_HUMAN 60S ribosomal protein L7 | HexNAc(1)Hex(1)NeuAc(1) | O | Cytoplasm | 3.03 |

NeuAc: N-acetylneuraminic acid; HexNAc: N-acetylhexosamine; Hex: hexose; Fuc: fucose.

**Supplementary Table S1. Continued**

| UniProt | Protein Name | Glycans | O/N | Location | Log Prob |
|---|---|---|---|---|---|
| P08236 | BGLR_HUMAN Beta-glucuronidase | HexNAc(3)Hex(3)Fuc(1)NeuAc(2) | O | Lysosome | 2.81 |
| Q9Y4L1 | HYOU1_HUMAN Hypoxia up-regulated protein 1 | HexNAc(3)Hex(3)Fuc(1)NeuAc(2) | O | Endoplasmic reticulum | 2.44 |
| Q96HR9 | REEP6_HUMAN Receptor expression-enhancing protein 6 | HexNAc(1)Hex(1)Fuc(1)NeuAc(1) | O | Endoplasmic reticulum membrane, cytoplasmic vesicle, clathrin-coated vesicle membrane | 2.34 |
| P11279 | LAMP1_HUMAN Lysosome-associated membrane glycoprotein 1 | HexNAc(4)Hex(4)Fuc(2)NeuAc(1) | O | Vacuole, membrane, endosome, cell surface | 2.06 |
| P49848 | TAF6_HUMAN Transcription initiation factor TFIID subunit 6 | HexNAc(2)Hex(2)Fuc(1)NeuAc(1) | O | Nucleus | 2.02 |
| Q14204 | DYHC1_HUMAN Cytoplasmic dynein 1 heavy chain 1 | HexNAc(1)Hex(1)NeuAc(2) | O | Cytoplasm, cytoskeleton | 1.95 |
| O00192 | ARVC_HUMAN Armadillo repeat protein deleted in velo-cardio-facial syndrome | HexNAc(2)Hex(1)Fuc(1)NeuAc(1) | O | Cell junction, adherens junction, nucleus, cytoplasm | 1.92 |
| O95299 | NDUAA_HUMAN NADH dehydrogenase [ubiquinone] 1 alpha subcomplex subunit 10, mitochondrial | HexNAc(3)Hex(2)Fuc(1)NeuAc(1) | O | Mitochondrion matrix | 1.9 |
| Q5T653 | RM02_HUMAN 39S ribosomal protein L2, mitochondrial | HexNAc(1)Hex(1)Fuc(1)NeuAc(1) | O | Mitochondrion | 1.72 |
| P78316 | NOP14_HUMAN Nucleolar protein 14 | HexNAc(1)NeuAc(1) | O | Nucleus, nucleolus | 1.67 |
| Q96HA1 | P121A_HUMAN Nuclear envelope pore membrane protein POM 121 | HexNAc(3)Hex(2)NeuAc(1) | O | Nucleus, nuclear pore complex, nucleus membrane, endoplasmic reticulum membrane | 1.66 |
| Q8IYB8 | SUV3_HUMAN ATP-dependent RNA helicase SUPV3L1, mitochondrial | HexNAc(2)Hex(1)NeuAc(1) | O | Nucleus, mitochondrion matrix, mitochondrion nucleoid | 1.57 |
| Q4KMP7 | TB10B_HUMAN TBC1 domain family member 10B | HexNAc(2)Hex(2)NeuAc(2) | O | Cytoplasm | 1.47 |

NeuAc: N-acetylneuraminic acid; HexNAc: N-acetylhexosamine; Hex: hexose; Fuc: fucose.

**Supplementary Table S1. Continued**

| UniProt | Protein Name | Glycans | O/N | Location | Log Prob |
|---|---|---|---|---|---|
| Q9UKM7 | MA1B1_HUMAN Endoplasmic reticulum mannosyl-oligosaccharide 1,2-alpha-mannosidase | HexNAc(1)Hex(1)NeuAc(2) | O | Endoplasmic reticulum membrane | 1.41 |
| Q9UHD1 | CHRD1_HUMAN Cysteine and histidine-rich domain-containing protein 1 | HexNAc(1)Hex(1)NeuAc(1) | O | Cytosol | 1.3 |

Glycopeptides were identified by Byonic following LC–MS/MS analysis. Protein accession, protein name, assigned glycan composition, glycosylation type (O- or N-linked), reported protein subcellular localisation, and Byonic log probability (Log Prob) are shown. NeuAc: N-acetylneuraminic acid; HexNAc: N-acetylhexosamine; Hex: hexose; Fuc: fucose.

**Supplementary Table S2. Glycopeptides with sialic acids in CMAH(+) LNCaP cells**

| UniProt | Protein Name | Glycans | O/N | Location | Log Prob |
|---|---|---|---|---|---|
| P00747 | PLMN_HUMAN Plasminogen | HexNAc(1)Hex(1)NeuAc(1) | O | Secreted | 16.26 |
| P02786 | TFR1_HUMAN Transferrin receptor protein 1 | HexNAc(1)Hex(1)NeuAc(1) | O | Cell membrane | 9.8 |
| P02786 | TFR1_HUMAN Transferrin receptor protein 1 | HexNAc(1)Hex(1)NeuAc(2) | O | Cell membrane, melanosome | 9.41 |
| P26572 | MGAT1_HUMAN Alpha-1,3-mannosyl-glycoprotein 2-beta-N-acetylglucosaminyltransferase | HexNAc(1)Hex(1)NeuAc(2) | O | Golgi apparatus membrane, cytoplasm, perinuclear region | 7.12 |
| Q9HC07 | TM165_HUMAN Transmembrane protein 165 | HexNAc(1)Hex(1)NeuAc(2) | O | Golgi apparatus membrane, golgi apparatus, lysosome membrane, early endosome membrane, late endosome membrane | 4.36 |
| Q8TD43 | TRPM4_HUMAN Transient receptor potential cation channel subfamily M member 4 | HexNAc(3)Hex(2)NeuAc(1) | O | Cell membrane, endoplasmic reticulum, golgi apparatus | 4.3 |
| P04004 | VTNC_HUMAN Vitronectin | HexNAc(4)Hex(5)NeuAc(2) | N | Secreted, extracellular space | 4.22 |
| P00734 | THRB_HUMAN Prothrombin | HexNAc(4)Hex(5)NeuAc(2) | N | Secreted, extracellular space | 4.05 |
| P00734 | THRB_HUMAN Prothrombin | HexNAc(3)Hex(4)NeuAc(1) | N | Secreted, extracellular space | 3.87 |
| P04114 | APOB_HUMAN Apolipoprotein B-100 | HexNAc(4)Hex(5)NeuAc(1) | N | Cytoplasm, secreted, lipid droplet | 3.55 |
| P00734 | THRB_HUMAN Prothrombin | HexNAc(4)Hex(5)NeuAc(1) | N | Secreted, extracellular space | 3.3 |
| P00734 | THRB_HUMAN Prothrombin | HexNAc(4)Hex(5)Fuc(2)NeuAc(1) | N | Secreted, extracellular space | 3.28 |
| Q96C36 | P5CR2_HUMAN Pyrroline-5-carboxylate reductase 2 | HexNAc(2)Hex(3)NeuAc(1) | O | Cytoplasm, mitochondrion | 3.26 |
| O15405 | TOX3_HUMAN TOX high mobility group box family member 3 | HexNAc(3)Hex(3)NeuAc(1) | O | Nucleus | 3.03 |
| Q8NHH1 | TTL11_HUMAN Tubulin polyglutamylase TTLL11 | HexNAc(3)Hex(3)NeuAc(2) | O | Cytoplasm, cytoskeleton, cilium basal body | 2.81 |
| Q52WX2 | SBK1_HUMAN Serine/threonine-protein kinase SBK1 | HexNAc(2)Hex(3)NeuAc(1) | O | Cytoplasm | 2.77 |

NeuAc: N-acetylneuraminic acid; NeuGc: N-glycolylneuraminic acid; HexNAc: N-acetylhexosamine; Hex: hexose; Fuc: fucose.

**Supplementary Table S2. Continued**

| UniProt | Protein Name | Glycans | O/N | Location | Log Prob |
|---|---|---|---|---|---|
| P49748 | ACADV_HUMAN Very long-chain specific acyl-CoA dehydrogenase, mitochondrial | HexNAc(3)Hex(3)NeuAc(1) | O | Mitochondrion inner membrane | 2.74 |
| Q9P203 | BTBD7_HUMAN BTB/POZ domain-containing protein 7 | HexNAc(3)Hex(1)Fuc(1)NeuGc(1) | O | Nucleus | 2.58 |
| Q96DX4 | RSPRY_HUMAN RING finger and SPRY domain-containing protein 1 | HexNAc(2)Hex(1)Fuc(2)NeuGc(1) | O | Secreted | 2.58 |
| A6NEQ2 | F181B_HUMAN Protein FAM181B | HexNAc(2)Hex(2)NeuAc(1) | O | Uncertain | 2.57 |
| Q8N2Y8 | RUSC2_HUMAN Iporin | HexNAc(1)NeuAc(1) | O | Cytoplasm, cytosol | 2.56 |
| Q7Z4W1 | DCXR_HUMAN L-xylulose reductase | HexNAc(3)Hex(3)Fuc(1)NeuAc(1) | O | Membrane | 2.4 |
| P12830 | CADH1_HUMAN Cadherin-1 | HexNAc(2)Hex(1)NeuAc(1) | O | Cell junction, adherens junction, cell membrane, single-pass type I membrane protein, endosome, golgi apparatus, trans-golgi network | 2.31 |
| P51610 | HCFC1_HUMAN Host cell factor 1 | HexNAc(2)Hex(1)NeuAc(1) | O | Cytoplasm, nucleus | 2.21 |
| Q96B97 | SH3K1_HUMAN SH3 domain-containing kinase-binding protein 1 | HexNAc(3)Hex(3)Fuc(1)NeuAc(1) | O | Cytoplasm, cytoskeleton, cytoplasmic vesicle membrane, peripheral membrane protein, synapse, synaptosome, cell junction, focal adhesion | 2.2 |
| P01871 | IGHM_HUMAN Immunoglobulin heavy constant mu | HexNAc(5)Hex(5)Fuc(1)NeuAc(1) | N | Secreted, cell membrane | 2.1 |
| P02743 | SAMP_HUMAN Serum amyloid P-component | HexNAc(4)Hex(5)NeuAc(2) | N | Secreted | 2.07 |
| Q9BTD8 | RBM42_HUMAN RNA-binding protein 42 | HexNAc(2)Hex(1)NeuAc(1) | O | Nucleus, cytoplasm | 2.02 |
| Q7RTV5 | AAED1_HUMAN Thioredoxin-like protein AAED1 | HexNAc(3)Hex(1) NeuAc(1) | O | Nucleus, nucleus speckle, cytoplasm, cytoplasmic granule | 2.01 |
| P49368 | TCPG_HUMAN T-complex protein 1 subunit gamma | HexNAc(1)Hex(1)NeuAc(2) | O | Cytoplasm | 1.9 |

NeuAc: N-acetylneuraminic acid; NeuGc: N-glycolylneuraminic acid; HexNAc: N-acetylhexosamine; Hex: hexose; Fuc: fucose.

**Supplementary Table S2. Continued**

| UniProt | Protein Name | Glycans | O/N | Location | Log Prob |
|---|---|---|---|---|---|
| P38432 | COIL_HUMAN Coilin | HexNAc(3)Hex(1)NeuAc(1) | O | Nucleus, cajal body | 1.84 |
| Q68EM7 | RHG17_HUMAN Rho GTPase-activating protein 17 | HexNAc(3)Hex(2)Fuc(1)NeuAc(1) | O | Cytoplasm, cell junction, tight junction | 1.63 |
| Q9UQ35 | SRRM2_HUMAN Serine/arginine repetitive matrix protein 2 | HexNAc(2)Hex(3)NeuAc(1) | O | Nucleus, nucleus speckle | 1.58 |
| P02751 | FINC_HUMAN Fibronectin | HexNAc(1)Hex(1)NeuAc(2) | O | Secreted, extracellular space, extracellular matrix | 1.52 |
| Q9H1B7 | I2BPL_HUMAN Interferon regulatory factor 2-binding protein-like | HexNAc(1)Hex(1)NeuAc(1) | O | Nucleus | 1.51 |
| Q92667 | AKAP1_HUMAN A-kinase anchor protein 1, mitochondrial | HexNAc(2)Hex(1)NeuAc(2) | O | Mitochondrion outer membrane, Mitochondrion | 1.49 |
| Q86VM9 | ZCH18_HUMAN Zinc finger CCCH domain-containing protein 18 | HexNAc(2)Hex(3)NeuAc(1) | O | Nucleus | 1.48 |
| P42858 | HD_HUMAN Huntingtin | HexNAc(1)Hex(1)NeuAc(3) | O | Cytoplasm, nucleus, early endosome | 1.48 |
| Q9ULI0 | ATD2B_HUMAN ATPase family AAA domain-containing protein 2B | HexNAc(1)Hex(1)Fuc(1)NeuGc(1) | O | Nucleus | 1.46 |
| O43824 | GTPB6_HUMAN Putative GTP-binding protein 6 | HexNAc(1)Hex(1)NeuGc(1) | O | Cytoplasm | 1.45 |
| P00734 | THRB_HUMAN Prothrombin | HexNAc(4)Hex(5)NeuAc(2) | N | Secreted, extracellular space | 1.44 |
| Q16643 | DREB_HUMAN Drebrin | HexNAc(2)Hex(1)NeuAc(1) | O | Cytoplasm, cell projection, cytoplasm, cell cortex, cell junction, cell projection, growth cone | 1.44 |
| Q9BWF3 | RBM4_HUMAN RNA-binding protein 4 | HexNAc(3)Hex(3)Fuc(1)NeuAc(1) | O | Nucleus, nucleus speckle, cytoplasm, cytoplasmic granule | 1.41 |

NeuAc: N-acetylneuraminic acid; NeuGc: N-glycolylneuraminic acid; HexNAc: N-acetylhexosamine; Hex: hexose; Fuc: fucose.

**Supplementary Table S2. Continued**

| UniProt | Protein Name | Glycans | O/N | Location | Log Prob |
|---|---|---|---|---|---|
| Q96KC8 | DNJC1_HUMAN DnaJ homolog subfamily C member 1 | HexNAc(3)Hex(1)NeuAc(1) | O | Endoplasmic reticulum membrane, nucleus membrane, microsome membrane | 1.4 |
| Q00796 | DHSO_HUMAN Sorbitol dehydrogenase | HexNAc(1)Hex(1)Fuc(1)NeuAc(1) | O | Mitochondrion membrane, peripheral membrane protein, cell projection, cilium, flagellum | 1.36 |
| P54886 | P5CS_HUMAN Delta-1-pyrroline-5-carboxylate synthase | HexNAc(1)Hex(1)NeuGc(1) | O | Mitochondrial inner membrane | 1.32 |
| P01871 | IGHM_HUMAN Immunoglobulin heavy constant mu | HexNAc(4)Hex(5)Fuc(1)NeuAc(1) | N | Secreted, cell membrane | 1.31 |

Glycopeptides were identified by Byonic following LC–MS/MS analysis. Protein accession, protein name, assigned glycan composition, glycosylation type (O- or N-linked), reported protein subcellular localisation, and Byonic log probability (Log Prob) are shown. NeuAc: N-acetylneuraminic acid; NeuGc: N-glycolylneuraminic acid; HexNAc: N-acetylhexosamine; Hex: hexose; Fuc: fucose.
